# The landscape of T cell infiltration in human cancer and its association with antigen presenting gene expression

**DOI:** 10.1101/025908

**Authors:** Yasin Şenbabaoğlu, Andrew G. Winer, Ron S. Gejman, Ming Liu, Augustin Luna, Irina Ostrovnaya, Nils Weinhold, William Lee, Samuel D. Kaffenberger, Ying Bei Chen, Martin H. Voss, Jonathan A. Coleman, Paul Russo, Victor E. Reuter, Timothy A. Chan, Emily H. Cheng, David A. Scheinberg, Ming O. Li, James J. Hsieh, Chris Sander, A. Ari Hakimi

## Abstract

**One sentence summary:** In silico decomposition of the immune microenvironment among common tumor types identified clear cell renal cell carcinoma as the most highly infiltrated by T-cells and further analysis of this tumor type revealed three distinct and clinically relevant clusters which were validated in an independent cohort.

**Abstract:** Infiltrating T cells in the tumor microenvironment have crucial roles in the competing processes of pro-tumor and anti-tumor immune response. However, the infiltration level of distinct T cell subsets and the signals that draw them into a tumor, such as the expression of antigen presenting machinery (APM) genes, remain poorly characterized across human cancers. Here, we define a novel mRNA-based T cell infiltration score (TIS) and profile infiltration levels in 19 tumor types. We find that clear cell renal cell carcinoma (ccRCC) is the highest for TIS and among the highest for the correlation between TIS and APM expression, despite a modest mutation burden. This finding is contrary to the expectation that immune infiltration and mutation burden are linked. To further characterize the immune infiltration in ccRCC, we use RNA-seq data to computationally infer the infiltration levels of 24 immune cell types in a discovery cohort of 415 ccRCC patients and validate our findings in an independent cohort of 101 ccRCC patients. We find three clusters of tumors that are primarily separated by levels of T cell infiltration and APM gene expression. In ccRCC, the levels of Th17 cells and the ratio of CD8^+^ T/Treg levels are associated with improved survival whereas the levels of Th2 cells and Tregs are associated with negative clinical outcome. Our analysis illustrates the utility of computational immune cell decomposition for solid tumors, and the potential of this method to guide clinical decision-making.

## Introduction

Tumors are complex environments, composed of transformed cells as well as stromal and immune infiltrates. Tumor-infiltrating cells can demonstrate either tumor-suppressive or tumor-promoting effects, depending on the cancer type or the tumor model. For instance, regulatory T cells (Tregs) and tumor associated macrophages (TAMs) have been associated with pro-tumor functions(1-3), whereas CD8^+^ T cells have been associated with improved clinical outcomes and response to immunotherapy(4-8). Antitumor activity of antigen-specific CD8^+^ T cells may underlie the efficacy of immune checkpoint blockade therapy(9-11) as such CD8^+^ T cells have been shown to increase in quantity and activity after treatment with these drugs.

CD8^+^ T cells are activated by peptide antigens presented on major histocompatibility class I (MHC-I) molecules. A CD8^+^ T cell can proliferate when its T cell receptor (TCR) recognizes antigens presented by MHC-I on a target cell, leading to an antigen-specific immune response that kills antigen bearing cells(12). All nucleated cells express antigen presenting machinery (APM) genes that code for MHC-I subunits and proteins necessary to process antigens and load them onto MHC-I. The APM genes can be upregulated by type II interferon (IFNγ), which is secreted by activated CD8^+^ T cells and other immune infiltrates. Upregulation of APM genes can lead to a cytotoxic feed-forward loop: more antigen presentation increases the number of T cells that find their cognate antigens, which in turn increases IFNγ release, antigen presentation and cytotoxicity. Yet, identification of CD8^+^ T cells alone is not sufficient to characterize the cytotoxic potential of the complex tumor microenvironment. The net inflammatory nature of the tumor can better be understood by quantifying the infiltration levels of diverse immune cell types.

Tumor immune infiltrates have largely been characterized by tissue-based approaches such as immunohistochemistry (IHC) and flow cytometry. These approaches are limited by a number of factors including the number of cell types that can be assayed simultaneously and the amount of tissue required. Computational techniques applied to gene expression profiles of bulk tumors can rapidly provide a broader perspective on the intratumoral immune landscape (13, 14).

ccRCC has been shown to be a highly immune-infiltrated tumor in multiple clinical and genomic studies(15, 16). A recent study found that transcript levels of two genes expressed by cytolytic cells (*GZMA* and *PRF1*) were highest in clear cell renal cell carcinoma (ccRCC) when compared to 17 other human cancers (13). The spontaneous regression seen in up to 1% of ccRCC cases is also thought to be largely immune-mediated(17). Additionally, ccRCC was historically one of the first malignancies to respond to immunotherapy, and continues to be among the most responsive (18-21). However, the mechanisms underlying high immune infiltration, spontaneous remissions and response to immunotherapy in this malignancy remain poorly understood.

The success of immune checkpoint blockade in melanoma and non-small cell lung cancer has largely been attributed to the high mutation burden in these tumors(10, 11). A higher number of tumor mutations is expected to result in greater numbers of MHC binding neo-antigens that have been proposed to drive tumor immune-infiltration and response to immunotherapy (10, 13, 22, 23). However, the modest mutation load of ccRCC compared with other immunotherapy-responsive tumor types(24) challenges the notion that neo-antigens alone can drive immune infiltration and response to immunotherapy in tumors.

As depicted in the workflow in Fig. 1a, we employed 24 immune cell type-specific gene signatures from Bindea *et al.*(14) (Fig. 1b) to computationally infer the infiltration levels in tumor samples (Step 1). We validated the gene signatures and our inference methodology using a ccRCC cohort from our institution (Step 2). We then defined a T cell infiltration score (TIS), an overall immune infiltration score (IIS) and an APM score to highlight the immune response differences between ccRCC(25) and 18 other tumor types profiled by The Cancer Genome Atlas (TCGA) research network (Step 3). Next, we characterized the immune-infiltration patterns in ccRCC patients by using the levels of 24 immune cells, angiogenesis, and expression of immunotherapeutic targets such as PD-1, PD-L1 and CTLA-4 (Step 4). We then investigated a suite of mechanisms that could potentially drive tumor immune-infiltration and explain the observed infiltration patterns in ccRCC. Finally, we validated our findings in an independent multi-platform ccRCC dataset(26) (Step 5). This integrative study utilizing rich whole-exome, whole-transcriptome, proteomic, and clinical data substantially improves our understanding of the tumor microenvironment in ccRCC and establishes an approach that can easily be extended to other human cancers.

**Fig. 1.**
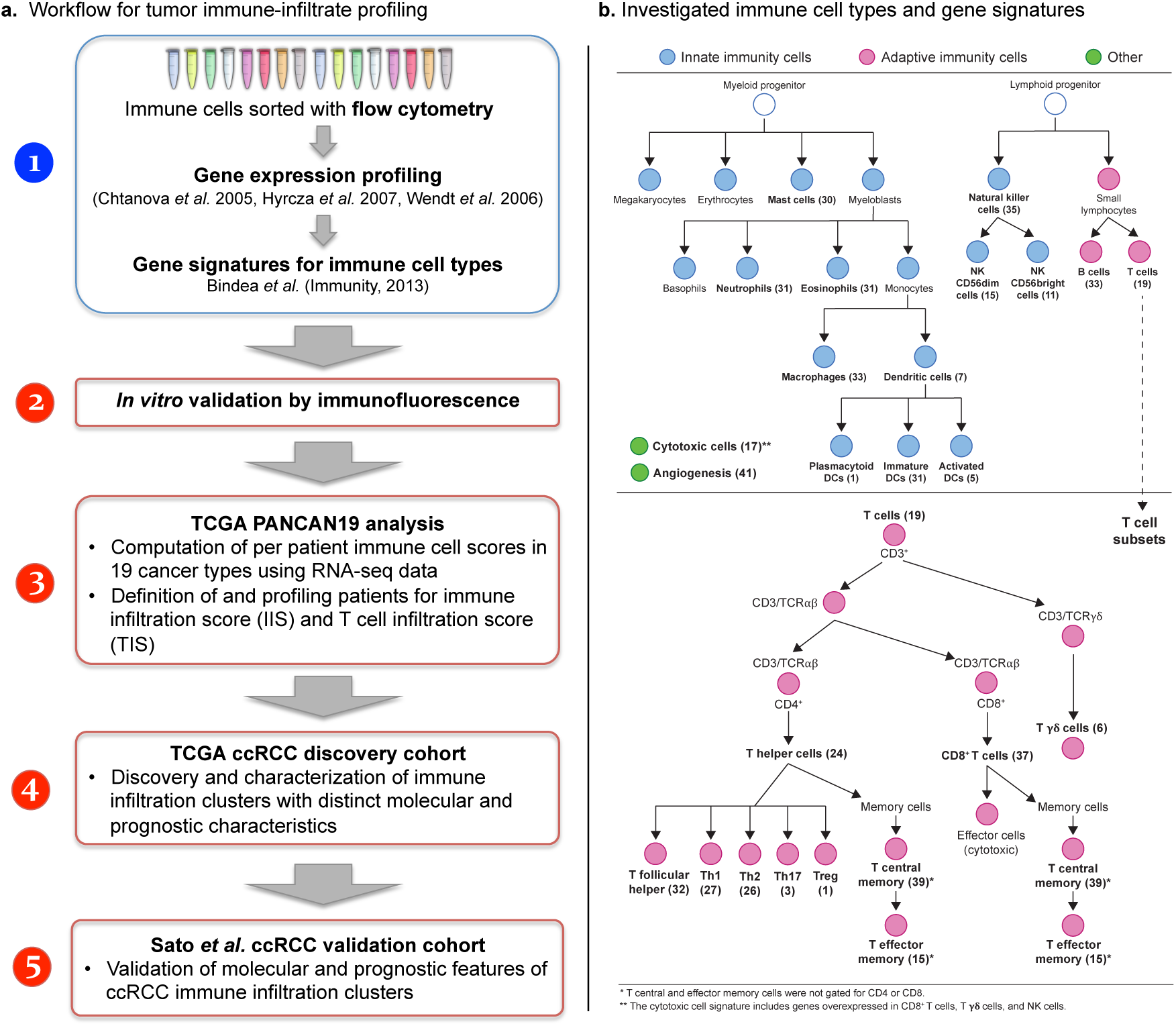
Workflow and the investigated immune cell types. **(a)** Workflow for tumor immune-infiltrate profiling. Gene signatures for 24 immune cell types were obtained from Bindea *et al.*(14) (Step 1). We computationally inferred the relative immune cell infiltration levels in cancer samples by computing an overrepresentation score (ssGSEA) from the transcript levels of the signature genes. We validated the immune cell scoring methodology by comparing ssGSEA scores with immunofluorescence staining and also with the transcript levels of immune cell markers (Step 2). Using this methodology, we profiled the immune infiltration levels in 19 human cancers, defined two novel immune infiltration scores, and showed the association of the T cell infiltration score (TIS) with antigen presenting gene expression across the tested cancer types (Step 3). We further characterized the immune infiltration patterns in clear cell renal cell carcinoma (ccRCC), the tumor type with the highest TIS median and one of the highest correlations between TIS and antigen presenting gene expression (Step 4). We investigated the association of ccRCC immune infiltration clusters with cancer-specific survival, neo-antigenicity, recurrent driver mutations, copy number alterations, and clinicopathologic variables. We validated our findings in an independent multi-platform ccRCC dataset(26) (Step 5). Red boxes denote analysis steps (2-5) while the blue box denotes the literature resources for gene signatures. **(b)** The investigated immune cell types are shown (**bold**) with two hierarchical trees: innate and adaptive immunity cell types (top panel), and distinct T cell subsets (bottom panel). The number of genes in each signature is displayed in parentheses next to the studied cell types.

## Results

## *In silico* decomposition and orthogonal validation of the tumor-immune microenvironment

We quantified the relative tumor infiltration levels of 24 immune cell types by interrogating expression levels of genes in published signature gene lists(14). The signatures we used comprised a diverse set of adaptive and innate immune cell types; and contained 509 genes in total (**Table S1**). 98.4% (501) of these genes were used uniquely in only one signature (Fig. S1). Due to the interconnectedness between immune cell infiltration and the antigen presenting machinery (APM), we also defined a 7-gene APM signature that consisted of MHC class I genes (HLA-A/B/C, B2M) and genes involved in processing and loading antigens (TAP1, TAP2 and TAPBP). mRNA-based scores for these signatures were then computed separately for each sample using single sample gene set enrichment analysis (ssGSEA)(27). ssGSEA measures the per sample overexpression level of a particular gene list by comparing the ranks of the genes in the gene list with those of all other genes.

We validated the gene signatures and the ssGSEA methodology in a series of internal and independent tests. We first performed an internal validation test for the immune cell gene signatures by using the three datasets(28-30) originally utilized by Bindea *et al.*(14) to derive the signatures. We asked whether the mRNA expression of the 501 unique signature genes had sufficient variance to discriminate between immune cell types separated by magnetic or fluorescence activated cell sorting. To this end, we obtained the microarray expression values for these genes, normalized with GCRMA(31) and corrected for batch effects using ComBat(32) (Fig. S2, **Materials and Methods**). We then computed the principal components (PC) of the batch-effect corrected dataset as a linear combination of the sorted immune cell types. This PC analysis successfully separated the cells into groups consistent with their hematopoietic lineage, suggesting adequate discrimination power for the signature genes (Fig. 2a). More specifically, PC1 and PC2 achieved the separation of the following four groups: 1) macrophages and dendritic cells, 2) B cells, NK cells (CD56dim and CD56 bright), CD8^+^ and CD4^+^ T cells, 3) Th1, Th2, T gamma delta and T follicular helper cells, 4) mast cells, neutrophils, and eosinophils. These 14 cell types only include those separated by magnetic or fluorescence activated cell sorting; *i.e.* exclude the ones for which signature genes are based on biological knowledge (such as Tregs and Th17 cells) or the ones that are umbrella terms (such as T helper cells and cytotoxic cells). The separation between CD8^+^ and CD4^+^ T cells was greatly enhanced if batch effect correction and PC analysis were performed with only the signatures genes of sorted T cell subpopulations (Fig. S3, Materials and Methods).

**Fig. 2.**
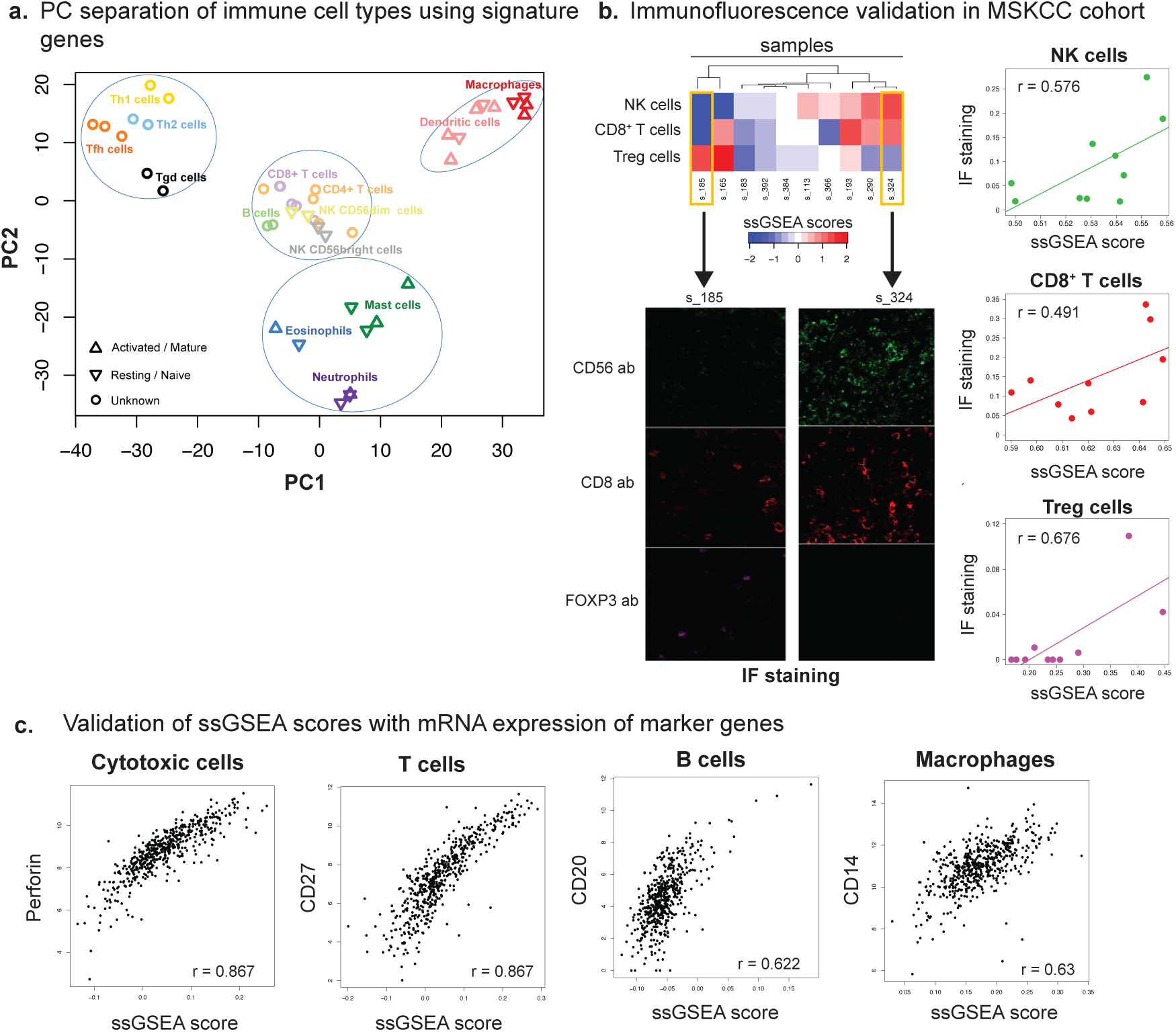
Validation of the immune cell scoring method. (**a)** Principal component (PC) analysis on immune cell types using transcript levels of signature genes. Microarray gene expression data were generated from immune cell types sorted by magnetic or fluorescent activated cell sorting(28-30) and used in Bindea *et al.*(14) to derive the signatures. **(b)** Immunofluorescence validation in MSKCC cohort. ssGSEA scores were computed using RNA-seq data to assess the infiltration levels of NK cells, CD8^+^ T cells, and Treg cells in 10 ccRCC tumors (top left panel). The infiltration levels of the same cell types were probed by immunofluorescence (IF) using the CD56, CD8 and FOXP3 antibodies respectively. IF staining is shown for two samples that are at opposite ends of the unsupervised hiearchy (bottom left panel). The association of the immune infiltrate levels inferred by these two orthogonal methods (ssGSEA and IF) is shown in the right panel where each dot represents a sample. Spearman correlations are given in the scatter plots. The IF staining level for a given sample was determined as the average across three representative regions on the slide. **(c)** Validation of ssGSEA scores with transcript levels of immune cell markers. The scatter plots show the scores for cytotoxic cells, T cells, B cells and macrophages in ccRCC plotted against the log2 RNA-seq values of marker genes: Perforin, CD267, CD20 and CD14 respectively. These genes are not found in the signatures used to compute ssGSEA scores. Spearman correlations are given in the scatter plots, and each point is a sample.

Next, we performed two independent tests to validate the joint quality of the signature genes and the ssGSEA methodology in inferring immune cell infiltration levels. The first validation test involved the comparison of mRNA-based ssGSEA scores with levels of immunofluorescence-stained immune cells from 10 MSKCC primary ccRCC tumors (**Materials and Methods** for sample preparation). Immunofluorescence (IF) staining was performed for three immune cell types that are extensively studied with immunohistochemistry: CD8^+^ T cells (anti-CD8 antibody), natural killer (NK) cells (anti-CD56 antibody) and Regulatory T cells (Tregs)(anti-FOXP3 antibody). We found that the quantitative estimates of immune cell infiltration from IF were well correlated with the ssGSEA scores (Fig. 2b). The Spearman correlation for the NK, CD8^+^ T, and Treg cell populations were 0.576, 0.491, and 0.676 respectively.

The second validation test involved the comparison of ssGSEA-based immune infiltration scores with the gene expression levels of key flow cytometry cell markers. We found that the ssGSEA scores for cytotoxic cells, T cells, B cells, and macrophages were highly correlated with the expression levels of the genes perforin, CD27, CD20, and CD14 respectively (Fig. 2c). We also observed moderate correlations for other cell types (Fig. S4, **Table S2**). These independent validation results provide further evidence that our *in silico* decomposition is a reliable and accurate method to infer immune infiltration levels in tumor samples.

## The T cell infiltration spectrum across 19 human cancers

We used the aforementioned 24 gene signatures to computationally assess the infiltration levels of immune cell types in 7567 tumor and 633 normal samples from 19 different cancer types profiled by TCGA (**Table S3**). To achieve a more focused view of the immune infiltration landscape in human cancers, we defined two aggregate scores: (1) the overall immune infiltration score (IIS) from both adaptive and innate immune cell scores, and (2) the T cell infiltration score (TIS) from nine T cell scores (CD8^+^ T, Th1, Th2, Th17, Treg, T effector memory, T central memory, T helper, and T cells) (**Materials and Methods**).

We next computed the TIS and IIS of each sample in the study as the sum of the relevant individual scores. Lung adenocarcinoma (LUAD) and ccRCC represented the highest end of the TIS and IIS spectrum (Fig. 3a, Fig. S5). Missense mutations within tumor cells are a known source of neo-antigens that can initiate a T cell dependent immune response(33). However, we did not observe a pan-cancer correlation between the immune infiltration levels of tumors and their respective number of somatic missense mutations (Fig. 3a for TIS, Fig. S5 for IIS, and Fig. S6 for a selected group of T cell subpopulations that make up the TIS). This result suggested that a factor other than mutation burden was necessary to explain the observed variation in immune infiltration levels. One notable exception was colorectal adenocarcinoma (COADREAD) where the hypermutated subpopulation had elevated levels of TIS (Pearson r=0.303, p=3.6×10^−7^).

**Fig. 3.**
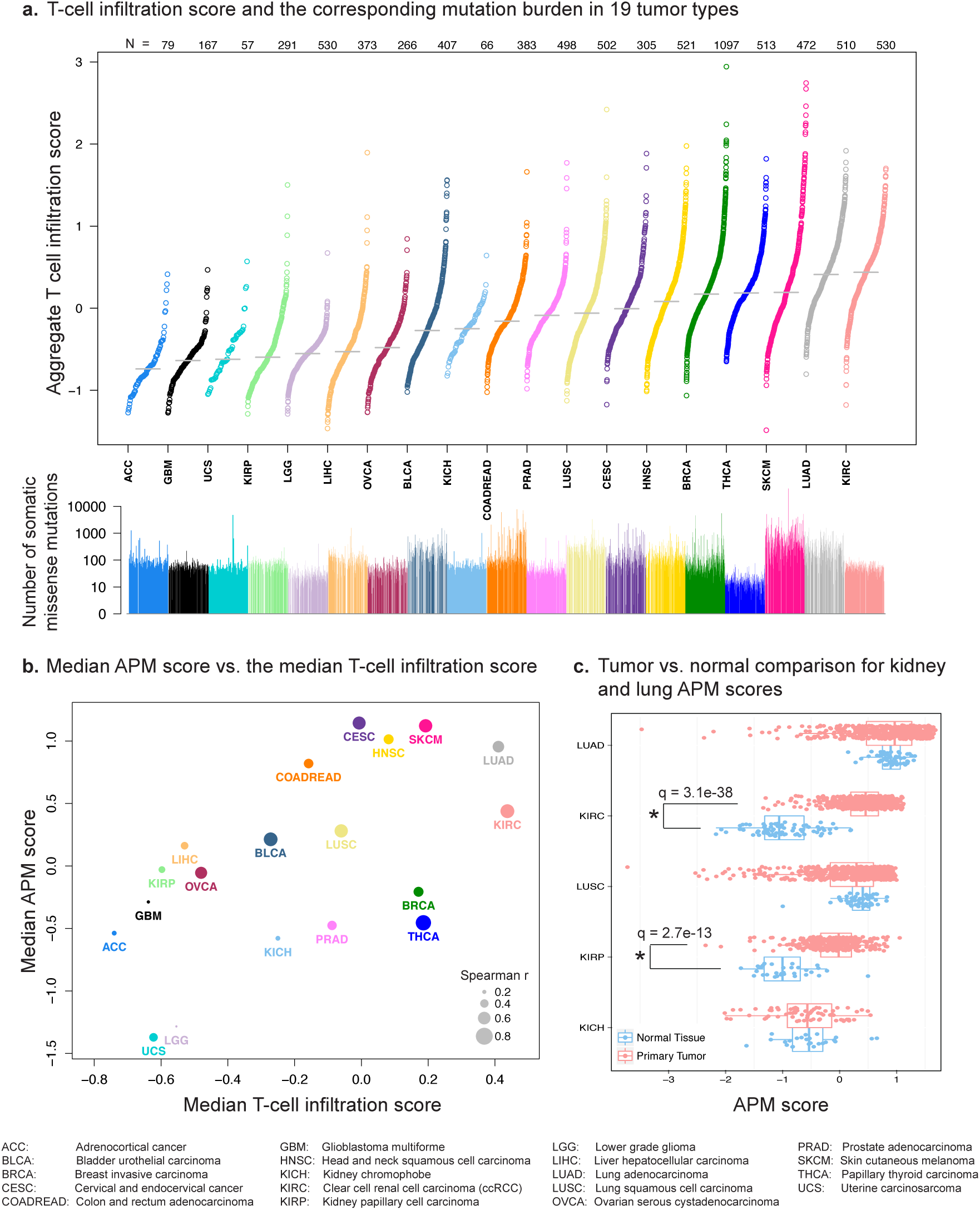
Pan-cancer analysis of T cell infiltration and its association with antigen presenting machinery (APM) gene expression. **(a)** T cell infiltration scores (TIS) and the corresponding mutation load in 19 tumor types. TIS is an aggregate score obtained as the average of nine distinct T cell subset scores (CD8^+^ T, Th1, Th2, Th17, Treg, T effector memory, T central memory, T helper, and T cells). Each circle in the top panel shows the TIS for a tumor sample. In the bottom panel, the vertical line corresponding to each circle shows the number of somatic missense mutations (log10 scale). Two immunotherapy-responsive tumors, clear cell renal cell carcinoma (KIRC) and lung adenocarcinoma (LUAD), have the highest TIS medians. KIRC has a modest mutation burden compared to LUAD. **(b)** The association between the median APM score and the median T cell infiltration score across 19 tumor types. The sizes of the circles are proportional to the within-cohort Spearman correlation between TIS score and APM score. KIRC and LUAD are among the highest not only for APM score but also for the APM–TIS correlation. **(c)** The APM score differences between tumors and adjacent normal tissue in kidney and lung neoplasms. Each circle is the APM score of a tumor (red) or an adjacent normal (blue) sample. No significant tumor–normal differences are observed in lung adenocarcinoma (LUAD), lung squamous cell carcinoma (LUSC), or kidney chromophobe (KICH) at α = 0.05. However, clear cell and papillary renal cell carcinoma (KIRC and KIRP) tumors significantly overexpress APM genes. The Benjamini-Hochberg adjusted p-values are reported in the figure (Mann-Whitney tests).

Immune infiltration is expected to increase the expression of APM genes in the tumor through paracrine signaling. Therefore, we investigated the correlation between the TIS and APM scores across the tested tumor types. As expected, the median TIS and the median APM score in the 19 cohorts showed a strong correlation (Spearman r=0.611), where ccRCC and LUAD were again among the highest with respect to the within-cohort TIS-APM correlation (Fig. 3b). Interestingly, a comparison of the APM expression between the tumor and normal tissue for kidney (clear cell, chromophobe and papillary sub-histologies) and non-small cell lung tumors (adenocarcinoma and squamous cell) revealed that the tumor-normal difference was highly significant for ccRCC (q=3.1×10^−38^, Mann-Whitney test) and papillary RCC (q=2.7×10^−13^, Mann-Whitney test) but not significant for other tumor types (Fig. 3c). APM expression of different stage and grade ccRCC tumors showed a lack of association between APM and either stage (p=0.263) or grade (p=0.118, Fisher’s exact tests) (Fig. S7 a,b). These results indicate that APM upregulation in ccRCC is likely a tumor-specific phenomenon caused by reasons other than necrosis. Melanoma (SKCM) is another immunotherapy-responsive tumor with high TIS (Fig. 3a), but data from adjacent normal tissue were not available for this tumor type.

In a survey of the other immune cell types, we found that the unique features of ccRCC immune infiltration extends to high levels of CD8^+^ T cells, plasmacytoid dendritic cells (pDC), T cells, cytotoxic cells, neutrophils; and low levels of Th2 and Treg cells compared with the other 18 cancer types (Fig. S8).

## Immune-infiltrate decomposition in ccRCC reveals three distinct patient clusters

In our effort to characterize the microenvironment of ccRCC tumors, we expanded our repertoire of 24 immune cell types to also include an angiogenesis signature(34) (**Table S1**), and immunotherapeutic targets PD-1 (*PDCD1*), PD-L1 (*CD274*) and CTLA-4 (*CTLA4*). Angiogenesis is well established to be a characteristic component of immune inflammation(35), and ccRCC is known to have high angiogenic capacity due to constitutive activation of the hypoxia-inducible factor pathway(36). We confirmed the high angiogenesis levels in ccRCC via a comparison against 18 other tumor types explored in this study (Fig. S8).

Using the ssGSEA scores from the expanded panel of 28 immune-and inflammation-related gene signatures, we performed unsupervised clustering on the TCGA cohort of 415 patients (**Materials and Methods**). Strikingly, this analysis revealed three distinct clusters that predominantly separated according to levels of T cell infiltration and APM gene expression, here termed the 1) T cell enriched, 2) heterogeneously infiltrated, and 3) non-infiltrated clusters (Fig. 4a). We observed that the T cell enriched tumors had markedly high expression of granzyme B (*GZMB*) and interferon-gamma (*IFNG*), effector molecules prominently associated with T cell response. An orthogonal measurement of purity by the DNA-based ABSOLUTE algorithm(37) confirmed that the T cell enriched and the non-infiltrated clusters were the least pure (mean 0.436) and the purest (mean 0.640) clusters respectively (p< 2×10^−16^, ANOVA). We then assessed the stromal content of samples using the RNA-based ESTIMATE algorithm(16) and investigated its association with the clusters. We found that the non-infiltrated cluster demonstrated the lowest stromal scores whereas the heterogeneous and T cell enriched clusters displayed mixed degrees of stromal content (p=4×10^−7^, ANOVA).

**Fig. 4.**
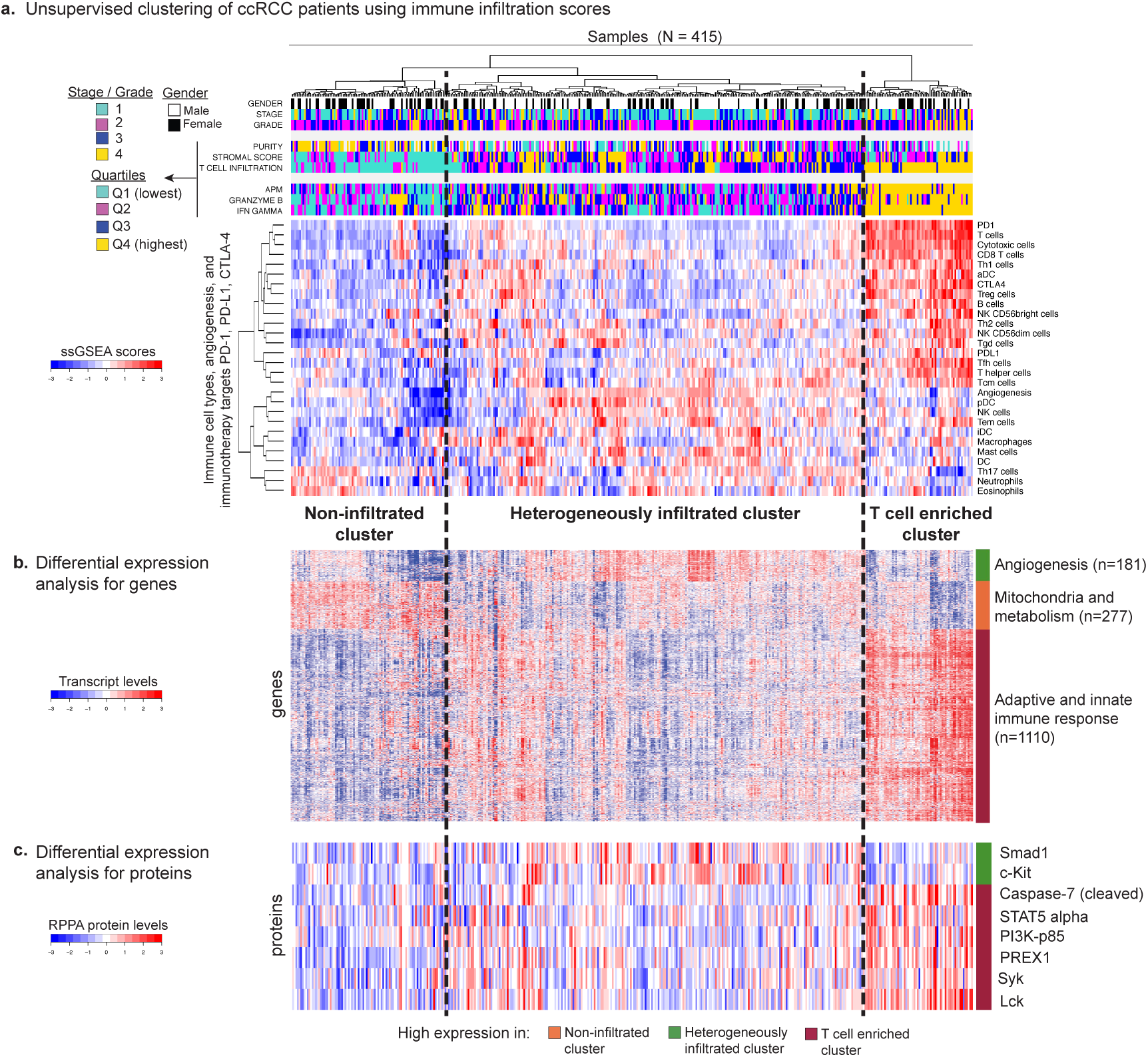
Characterization of immune infiltration clusters in ccRCC. **(a)** Unsupervised clustering of 415 ccRCC patients from the TCGA cohort using ssGSEA scores from 24 immune cell types, 3 immunotherapy targets (PD-1, PD-L1, CTLA-4), and angiogenesis. Hierarchical clustering was performed with Euclidean distance and Ward linkage. We discover three distinct immune infiltration clusters, here termed 1) non-infiltrated, 2) heterogeneously infiltrated, and 3) T cell enriched. The T cell enriched cluster is characterized by tumors with high APM scores and high granzyme B and interferon gamma mRNA expression levels. **(b)** Differential expression analysis with Mann-Whitney tests for all genes in the TCGA RNA-seq dataset excluding signature genes. Only genes that are significantly overexpressed in one cluster at a q-value cutoff of 5×10^−5^ are shown. Pathway analysis using DAVID(38) reveals that the genes overexpressed in the three clusters (N=1110, 181, and 277 respectively) are enriched in 1) adaptive and innate immune response, 2) angiogenesis, and 3) mitochondrial and metabolic processes. **(c)** Differential expression analysis with Mann-Whitney tests for all proteins in the TCGA reverse phase protein array (RPPA) dataset. Only proteins that are significantly overexpressed in one cluster at a q-value cutoff of 0.01 are shown. This analysis recapitulates the significant differences in immune response in the T cell enriched cluster, and in angiogenesis in the heterogeneously infiltrated cluster.

In order to validate the three immune infiltration clusters, we utilized a separate publicly available dataset of 101 ccRCC tumors for which comparable multi-platform data were available(26), and refer to it as the SATO dataset from here on. A random-forest classifier trained on the TCGA cohort was used to predict the immune infiltration class for each SATO patient (**Materials and Methods**). The heatmap of the same 28 immune features in the SATO dataset confirmed the existence of the three classes as well as elevated expression levels of APM, granzyme B and interferon-gamma in the T cell enriched cluster (Fig. S9a).

To further characterize the clusters’ unique molecular features, we next performed an unbiased analysis of differential gene and protein expression between the clusters. We excluded the signature genes and performed pathway analysis(38) for the genes significantly overexpressed in one of the clusters (q < 5×10^−5^, Mann-Whitney test). We observed that the T cell enriched group had significant overexpression of both adaptive and innate immunity genes (Fig. 4b and **Table S4**). On the other hand, the non-infiltrated group had significant overexpression of metabolism-and mitochondria-related genes (**Table S5**), while the heterogeneously infiltrated group had overexpression of angiogenesis-related genes (**Table S6**) (q < 5×10^−5^, Mann-Whitney test). These findings were again validated in the SATO dataset (Fig. S9b, Table S7a-7c). We next utilized the TCGA reverse phase protein array (RPPA) dataset for the differential protein expression analysis. We consistently observed overexpression of immune-related proteins, such as Lck and Syk, for the T cell enriched group; and an overexpression of angiogenesis related proteins, such as Smad1 (39, 40) and c-Kit (41-43), for the heterogeneously infiltrated group (q < 0.01, Mann-Whitney tests) (Fig. 4c). A proteomic dataset for the SATO cohort was not available.

We observed in Fig. 4a and 4b that the T cell enriched cluster had two subclusters, here termed TCa and TCb (Fig. 5a), with different immune cell infiltration and gene expression profiles. Gene set enrichment analysis with DAVID(38) and ClueGO(44) revealed that the genes overexpressed in TCa (q<5×10^−5^, Mann-Whitney test) were associated with metabolic and mitochondrial processes (Fig. 5b, Table S8a). The genes overexpressed in TCb (q<5×10^−5^, Mann-Whitney test) were enriched for processes related to cell cycle, extracellular matrix (ECM), and cellular proliferation (Fig. 5b, **Table S8b)**. We also found that these two subclusters had prognostic differences (Fig. 5c), with the TCb patients having worse cancer-specific survival than the TCa patients (p=0.0162, log-rank test). Moreover, the TCb subcluster had significantly higher macrophage infiltration (p=5.7×10^−4^) and stromal score (p=4.6×10^−4^, Mann-Whitney tests) with a moderate correlation between these two variables (Spearman r=0.418, p=5.8×10^−4^). This correlation generalized to the entire cohort (Spearman r=0.561, p<2×10^−16^), suggesting the possibility of macrophage recruitment by stromal cells(45) (Fig. S10). These results confirm the biologically distinct characteristics of the TCa and TCb subclusters within the T cell enriched group.

**Fig. 5.**
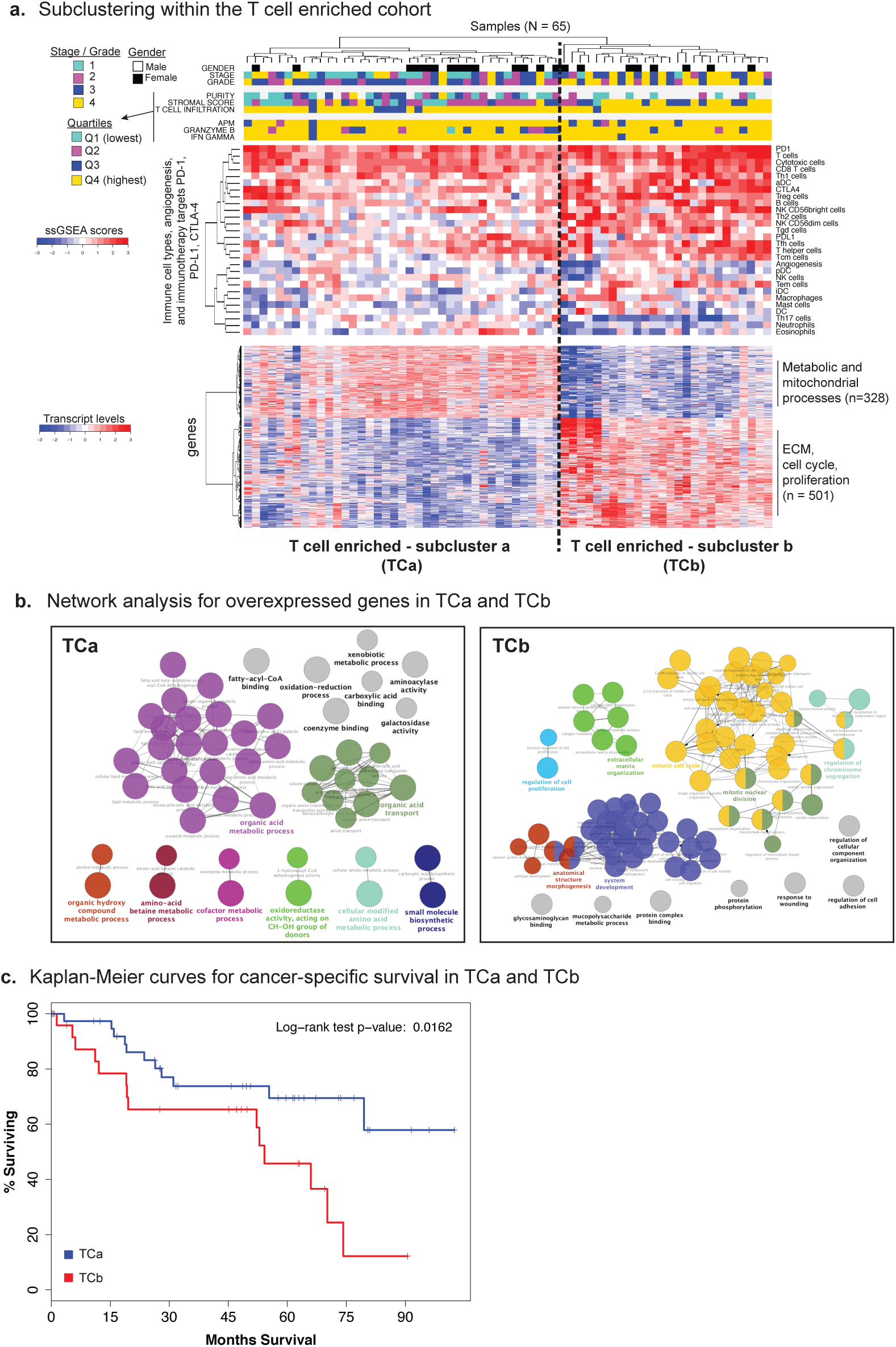
Subclustering within the T cell enriched cohort demonstrates gene expression and survival differences. **(a)** Hierarchical clustering within the T cell enriched cohort revealed two distinct subclusters, here termed TCa and TCb, that had differences in immune cell levels such as macrophages as well as in grade, stage, and stromal score (top panel). Hierarchical clustering was performed with Euclidean distance and Ward linkage. Differential gene expression analysis was performed with Mann-Whitney tests (bottom panel). Only genes that are significantly overexpressed in one cluster at a q-value cutoff of 5×10^−5^ are shown. Pathway analysis using DAVID(38) reveals that the genes overexpressed in TCa and TCb (N = 328 and 501 respectively) are enriched in 1) metabolic and mitochondrial processes; 2) and extracellular matrix (ECM), cell cycle and cell proliferation respectively. **b)** Network analysis with ClueGO(44) highlights the upregulation of metabolic processes in TCa, and the upregulation of ECM, cell cycle, cell proliferation in TCb. **(c)** Kaplan-Meier curves for cancer-specific survival in the TCa and TCb patients. Patients in the TCa subcluster have significantly better survival (log-rank test p-value = 0.016)

We next investigated whether the immune infiltration classes predicted by our mRNA-based decomposition algorithm were robust to intratumoral heterogeneity. We obtained a microarray gene expression dataset from the Gerlinger *et al.* (46) ccRCC multiregion tumor study (referred to as GERLINGER from here on). This dataset includes 56 tumor and 6 normal samples from 9 ccRCC patients. The authors sampled several tumor regions from each patient to investigate intratumor heterogeneity. We applied the random forest classifier trained on the TCGA ccRCC cohort to the GERLINGER samples to predict their immune infiltration class (Fig. S11). Interestingly, different regions from the same tumor showed very similar immune infiltration patterns if the tumor had a strong T cell enriched phenotype. Other tumors showed intratumor differences in terms of immune infiltration patterns **(Fig. S11)**.

## T cell infiltration levels are associated with clinical outcomes

We found that tumor immune-infiltration in ccRCC was associated with distinct clinicopathologic features. Male patients (p=0.018), higher stage (p=0.006) and higher grade (p=0.003) tumors were overrepresented in the T cell enriched class compared to the non-and-heterogeneously infiltrated groups (Fisher’s exact tests). Patients in the T cell enriched class had the poorest cancer-specific survival whereas the non-infiltrated group fared the best (p=0.05; log-rank test) (Fig. 6a).

**Fig. 6.**
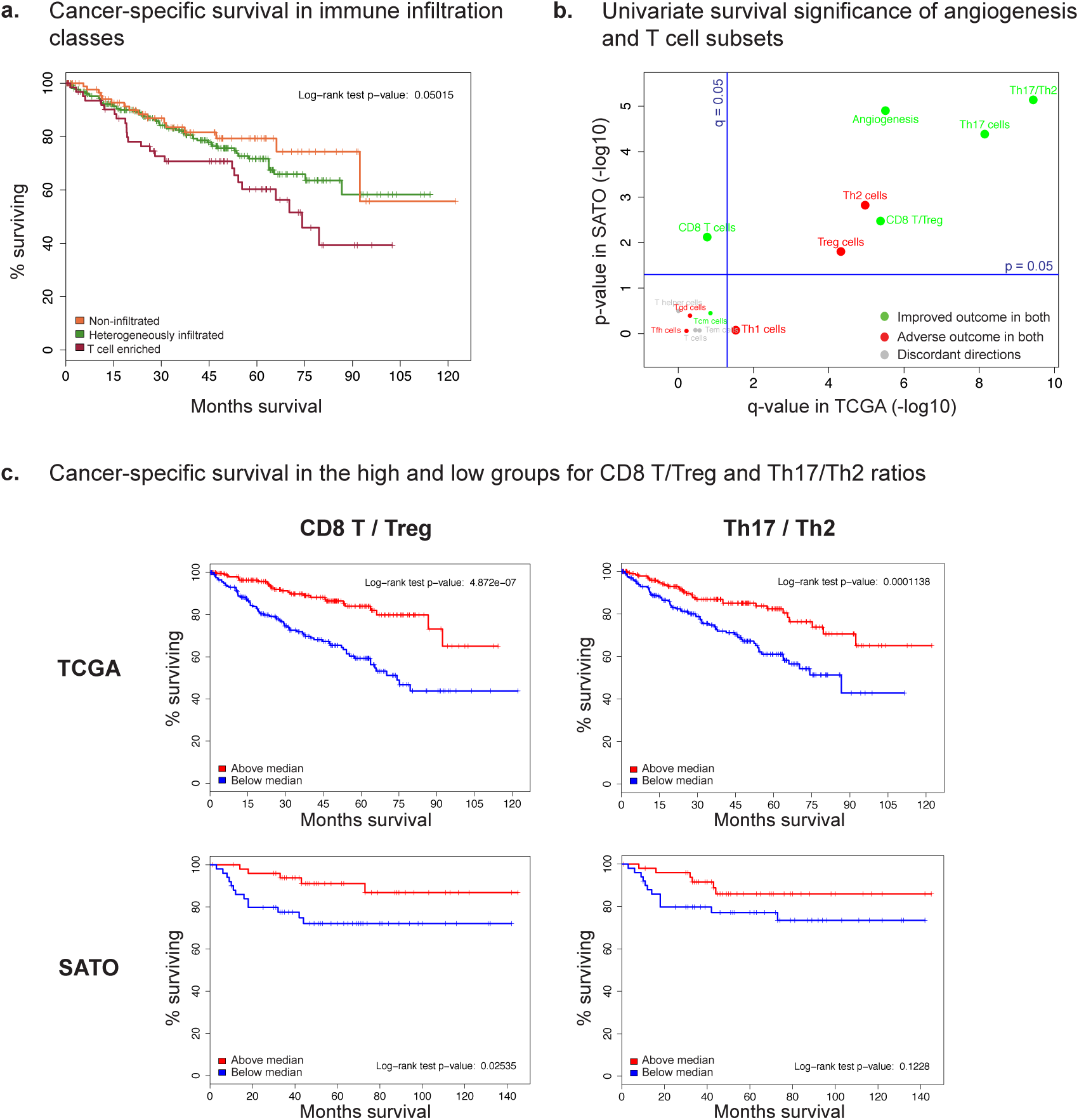
Prognostic significance of ccRCC immune infiltration classes and distinct T cell subsets. **(a)** Kaplan-Meier curves for cancer-specific survival in ccRCC immune infiltration classes. The T cell enriched class has the poorest survival whereas the non-infiltrated class is associated with better outcomes (log-rank test p-value = 0.05) Prognostic significance of angiogenesis and distinct T cell subsets in ccRCC. Univariate Cox proportional-hazards was used to regress ssGSEA scores on cancer-specific survival. The resultant p-values in the TCGA dataset were adjusted for multiple hypothesis testing, log-transformed, and then plotted against the log-transformed p-values from the SATO dataset. Survival associations concordant in both datasets are denoted in green and red for improved and poor outcome respectively. Discordant associations are denoted in grey. Th17 is the most significant pro-survival cell type in both TCGA and SATO datasets. However, the Th17/Th2 ratio is more significantly associated with improved survival compared with Th17 alone. In TCGA, CD8^+^ T cells are not significantly associated with improved survival at α = 0.05, but the CD8^+^ T/Treg ratio is. **(c)** Kaplan-Meier curves for cancer-specific survival in the high and low groups for the CD8^+^ T/Treg and Th17/Th2 ratios. Treg cells may inhibit CD8^+^ T cells, reducing their cytotoxic potential. Th17 and Th2 cells have both highly significant but opposite associations with cancer-specific survival. The median values for these two ratios are able to stratify both the TCGA and the SATO cohorts into groups with significant survival differences.

We then investigated the univariate significance of each T cell subset and angiogenesis as a predictor of cancer-specific survival. Cox proportional-hazards regression showed, in both the TCGA and SATO datasets, that the levels of Th17 cells and angiogenesis were strongly associated with favorable outcomes, whereas Th2 and Treg cells were associated with adverse outcomes (Fig. 6b) consistent with previous reports(15, 34, 47-50). The survival significance of the Th17/Th2 ratio surpassed those of Th17 and Th2 levels alone. Moreover, we observed that CD8^+^ T cell levels alone were not significantly associated with improved survival in the TCGA cohort, but the CD8^+^ T/Treg ratio was (Fig. 6b-c). Additional analyses demonstrated that previously identified prognostic features such as tumor stage and molecular subtype (ccA/ccB)(51) were associated with similarly prognostic immune infiltration scores. In particular, Treg and Th17 infiltration levels had respectively negative and positive association with tumor stage (q=1.2×10^−6^, ANOVA) (Fig. S12a). Treg and Th2 infiltration levels were higher in ccB subtype tumors, which have poor prognosis relative to ccA (q=3.9×10^−9^ and 1.2×10^−8^, Mann-Whitney tests) (Fig. S12b). In contrast, Th17 and CD8^+^ T cell infiltration levels were higher in ccA tumors (q=2.8×10^−12^ and 5.8×10^−6^, Mann-Whitney tests).

## Lack of association with immune infiltration, genomic alterations and neo-antigens

In light of our evidence suggesting the presence of immunologically distinct subsets of ccRCC tumors, we investigated mutation load and recurrent genomic alterations as potential drivers of the observed T cell infiltration. The tumors from the non-infiltrated class harbored slightly more somatic missense mutations than the T-cell-enriched class (the median number of somatic missense mutations in the non-infiltrated group was 36.5 versus 33 in the T cell enriched group; q=0.07). Out of the 11 driver genes commonly mutated in ccRCC, only *PBRM1* was mutated at significantly different rates between the three populations (**Fig.S13a**; higher in non-vs. T cell enriched q=0.04; higher in heterogeneous vs. T cell enriched; q=0.04). However, this observation was not validated in the SATO dataset. None of the common arm-level CNVs observed in ccRCC tumors were found at different rates between the three groups (Fig. S13b). Cancer neo-antigens have been demonstrated to drive T cell infiltration of tumors in murine models of cancer(33, 52). We hypothesized that the abundance or quality of cancer neo-antigens might differ between our tumor classes. To address this theory, we determined the HLA-A, HLA-B and HLA-C alleles of each ccRCC TCGA patient using OptiType(53). We then predicted the protein alterations expected to result from missense mutations in each tumor and identified those predicted to bind to MHC-I molecules (**Materials and Methods**). We found no significant difference in the median MHC-I binding count (Fig. S13c) or median binding affinity (Fig. S13d) of neo-antigens between the three classes of TCGA tumors. We also found no significant difference in the fraction of tumors with non-silent somatic mutations in an expanded set of APM genes (**Table S9a-c**). These results suggest that factors other than genomic alterations may be contributing to the immune infiltration of ccRCC tumors.

## Discussion

In this pan-cancer analysis, we present a novel approach for profiling the immune infiltration patterns of tumors using an mRNA-based computational decomposition method. Our data highlighted the immunotherapy-responsive tumors ccRCC and LUAD as having the highest T cell infiltration median. Moreover, ccRCC, but not LUAD, demonstrated significant upregulation of antigen presentation machinery in comparison with adjacent normal tissue.

Preliminary evidence emerging from clinical trials of immune checkpoint blockade therapy suggests that high mutation burdens may be predictive of good responses in NSCLC and melanoma(10, 11). However, ccRCC is another immunotherapy-responsive tumor despite bearing orders of magnitudes fewer mutations than NSCLC and melanoma. We suggest that ccRCC tumors may be responsive to checkpoint blockade because of a potent pre-existing immune infiltration and overall elevated level of antigen presentation and recognition.

Unsupervised clustering of ccRCC tumors using immune infiltration levels revealed three clusters of differentially infiltrated tumors, which were subsequently validated in an independent cohort. In particular, we found that the T cell enriched cluster was characterized by high expression levels of immune-response related genes including the immune checkpoint genes PD-1, PD-L1, and CTLA-4. Interestingly, a recent study also identified an aggressive, sunitinib resistant molecular subtype of metastatic ccRCC with cellular and molecular characteristics similar to the T cell enriched tumors discovered here(54). These findings across several cohorts of ccRCC patients suggest that a subset of ccRCC tumors may be both highly immune-infiltrated and immunosuppressed, as indicated by elevated expression of immune-checkpoint surface markers.

Our in-depth analysis including driver mutations, CNVs, mutation burden and neo-antigens failed to reveal any molecular mechanisms for the differential immune infiltration in ccRCC clusters. However, the lack of association between immune infiltration and predicted MHC-I binding tumor neo-antigens does not rule out neo-antigens as drivers of immune infiltration. Computational techniques for the prediction of immunogenic neo-antigens are not yet mature: most studies that identify immunogenic epitopes rely on a combination of computational, biochemical and cellular techniques. Overall, our results suggest that genetic alterations, mutation burden and predicted neo-antigens currently provide an incomplete explanation for the degree of immune infiltration in ccRCC.

Our findings underscore the prognostic significance of specific T cell subsets, consistent with previous tissue-based studies of ccRCC and other tumor types(55). Our results also illustrate the utility of ssGSEA for inferring immune infiltration levels in tumor specimens. The methodology in this study could directly be extended to the investigation of immune infiltration and its potential drivers in other tumor types and in various clinical settings. One example is colorectal cancer where we observe an intriguing association between the T cell infiltration and hypermutated samples. Ultimately, our approach enables the determination of a diverse array of immune infiltration patterns from small amounts of tissue such as biopsy samples; a strategy which could easily be incorporated into the clinical and trial setting.

## Materials and Methods

### Gene signatures and scoring for infiltration/activity levels with ssGSEA

Marker genes for immune cell types were obtained from Bindea *et al.*(14). Angiogenesis marker genes were obtained from Masiero *et al* (34). A signature of antigen presentation was created based on genes involved in processing and presentation of antigens on MHC(12). All signature genes are listed in **Table S1.** Infiltration levels for immune cell types, and activity levels for angiogenesis and antigen presentation were quantified using **ssGSEA**(27) in the R package **gsva**(56). **ssGSEA** takes as input the genome-wide transcriptional profile of a sample, and computes an overexpression measure for a gene list of interest relative to all other genes in the genome.

### Internal validation of gene signatures using principal component (PC) analysis

We performed an internal validation of the immune cell gene signatures on the three HG-U133A microarray datasets(28-30) originally used by Bindea *et al.*(14) to derive the signatures. The combined dataset had a total of 46 samples from 14 unique immune cell types. We first performed background correction and quantile normalization on the CEL files using GCRMA(31). We then performed two consecutive PC analysis to investigate the separation of (1) all 14 immune cell types, and (2) only the T cell subpopulations among the set of 14 cell types.

1. **PC separation of all immune cell types**: We reduced the GCRMA-normalized dataset to the signature genes by mapping the Affymetrix U133A probeset identifiers to HGNC symbols with the R biomaRt package(57), and filtering out the zero variance probesets. 840 probesets remained, corresponding to the 501 unique genes used in the immune cell signatures. A PC analysis on the normalized and reduced dataset revealed batch effects from the three data sources (Fig. S2, top panel). We corrected for batch effects using the nonparametric option in ComBat(32) (Fig. S2, bottom panel), and subsequently performed PC analysis on the 46 samples to investigate the separation of immune cell types by the first two PCs (Fig. 2a).
2. **PC separation of 6 T cell subpopulations**: We reduced the GCRMA-normalized dataset to the 19 T cell subpopulation samples and only the T cell related signature genes in a similar manner as (1). 400 probesets remained, corresponding to the 225 unique T cell subpopulation signature genes. Batch effects were corrected using the nonparametric option in ComBat(32), and PC analysis was subsequently performed on the 19 samples to investigate the separation of T cell subpopulations (Fig. S3).

### Multiplex immunofluorescence staining

Informed consent for tissue analysis was obtained under institutional review board-approved protocol IRB #89-076. Unstained pathologic slides of ten renal tumors from previously untreated patients who underwent either radical or partial nephrectomy for sporadic, resectable ccRCC were obtained and reviewed by a genitourinary pathologist. Paraffin-embedded tissue sections were de-waxed with xylene and rehydrated by gradient ethanol solutions. Antigen retrieval was then performed and the sections were subsequently blocked by bovine serum albumin plus serum with the addition of mouse monoclonal anti-human CD8 (Dako, clone C8/144B, catalogue #M7103 (58)), CD56 (Thermo scientific, clone 56C04, catalogue #MS-1149-P1 (59)) and FOXP3 (Abcam, clone 236A/E7, catalogue #ab20034 (60)). The sections were incubated with HRP-conjugated anti-mouse antibodies. TSA plus kits (Perkin Elmer) were used according to the manufacturer’s instructions. Finally, Leica upright confocal microscope was used to capture images. In order to quantify the degree of cellular infiltration, the individual positive cells for CD56, CD8 and FOXP3 were counted in three representative regions of each tumor. The ratio of CD56, CD8 and FOXP3 positive cells versus total cells (DAPI-stained) were determined.

### Gene and protein expression datasets

The pancan normalized gene-level RNA-seq data for the TCGA cohorts were downloaded from the UC Santa Cruz Cancer Genomics Browser(61) (https://genome-cancer.ucsc.edu/). These cohorts consisted of adrenocortical cancer (ACC), bladder urothelial carcinoma (BLCA), lower grade glioma (LGG), breast invasise carcinoma (BRCA), cervical and endocervical cancer (CESC), colon and rectum adenocarcinoma (COADREAD), glioblastoma multiforme (GBM), head and neck squamous cell carcinoma (HNSC), kidney chromophobe (KICH), kidney clear cell carcinoma (KIRC), kidney papillary cell carcinoma (KIRP), liver hepatocellular carcinoma (LIHC), lung adenocarcinoma (LUAD), lung squamous cell carcinoma (LUSC), ovarian serous cystadenocarcinoma (OVCA), prostate adenocarcinoma (PRAD), skin cutaneous melanoma (SKCM), thyroid carcinoma (THCA), and uterine carcinosarcoma (UCS).

TCGA ccRCC-specific analyses were performed with the KIRC datasets downloaded from Firebrowse (http://firebrowse.org). RSEM-normalized gene level data and reverse phase protein array (RPPA) data were used for gene and protein expression analyses respectively. Samples that had RNA-seq, mutation, and clinical data (N=415) were included in the discovery phase of the immune infiltration clusters.

The Sato *et al.*(26) Agilent microarray gene expression dataset was downloaded from ArrayExpress (http://www.ebi.ac.uk/arrayexpress/experiments/E-MTAB-1980/); and all samples (N=101) were included in the analysis. The probe identifiers in the Agilent platform were mapped to HGNC gene symbols, and the arithmetic mean across identifiers was used for cases where multiple Agilent identifiers mapped to a single HGNC symbol.

The Gerlinger *et al.* (46) Affymetrix Human Gene 1.0 ST microarray gene expression dataset was obtained via personal communication with the authors on November 10^th^, 2014. This dataset includes 56 tumor and 6 normal samples from 9 ccRCC patients. All samples were included in our analysis. The probe sets in this Affymetrix platform were mapped to HGNC gene symbols, and the geometric mean across probe sets was used for cases where multiple probe sets mapped to a single HGNC symbol.

The MSKCC RNA-seq dataset was generated and pre-processed internally. Raw output BAMs were converted back to FASTQ using PICARD Sam2Fastq. Maps were then mapped to the human genome using STAR aligner(62). The genome used was HG19 with junctions from ENSEMBL (GRCh37.69_ENSEMBL) and a read overhang of 49. Then any unmapped reads were mapped to HG19 using BWA MEM (version 0.7.5a). The two mapped BAMs were then merged and sorted and gene level counts were computed using htseq-count (options -s y -m intersection-strict) and the same gene models as used in the mapping step.

### TCGA PANCAN mutation calls

PANCAN mutation calls were downloaded from the BROAD Firehose’s stddata_2015_02_04 dataset (http://gdac.broadinstitute.org/). Additional COADREAD mutation calls were obtained from the MSKCC cBio portal(63) via personal communication. These mutation calls were used for all analyses, excluding neo-antigen analysis.

### Clinical data for TCGA and SATO patients

Clinical data for the TCGA dataset were obtained from the supplementary files of the ccRCC marker paper(25) (KIRC+Clinical+Data+Jul-31-2012). Vital status was determined from the field “Composite Vital status". Clinical data for the Sato dataset were obtained through direct communication with the authors.

### T cell infiltration and immune infiltration scores

The ssGSEA scores for each individual immune cell type were standardized across all tumor and normal samples in the investigated 19 tumor types (N = 8200). The T cell infiltration score was defined as the mean of the standardized values for all T cell subsets except for T gamma delta and T follicular helper cells: CD8 T, T helper, T, T central and effector memory, Th1, Th2, Th17, and Treg cells. T gamma delta and T follicular helper cells were excluded from the aggregate scores because it has been reported by the authors of one of the microarray datasets used by Bindea *et al*.(14) that some T cell specific genes were expressed in healthy brain tissue(28). Tissue gene expression maps verify that C1orf61 and FEZ1 in the T gamma delta signature; and B3GAT1, HEY1, CHGB and CDK5R1 in the T follicular helper signature are expressed at elevated levels in healthy brain tissue relative to other tissues(64).

The overall immune infiltration score for a sample was similarly defined as the mean of the standardized values for macrophages, dendritic cell (DC) subsets (total, plasmacytoid, immature, activated), B cells, cytotoxic cells, eosinophils, mast cells, neutrophils, NK cell subsets (total, CD56bright, CD56dim), and all T cell subsets excluding T gamma delta and T follicular helper cells.

### HLA typing and HLA-binding neoepitope prediction

Whole exome sequences for the TCGA KIRC tumors were downloaded using cgquery (https://cghub.ucsc.edu/). Whole-exome sequences for the SATO dataset were downloaded from the European Genome-phenome Archive (https://www.ebi.ac.uk/ega/studies/EGAS00001000509). BAM files containing whole exome sequences from normal and/or tumor samples were processed to obtain fastq files. Reads that aligned to HLA-A, HLA-B or HLA-C genes using RazerS3(65) (http://www.seqan.de/projects/razers/) were passed as input to OptiType v1.0(53) (https://github.com/FRED-2/OptiType). Discrepancies in HLA typing were resolved by consensus or exclusion. A MAF files containing missense mutations for each TCGA patient was obtained from cBioPortal (http://www.cbioportal.org/). A MAF file containing missense mutations for each SATO patient was obtained from the publication(26). Samtools (v 0.1.19) and snpEff (v3.5c) were used to identify the protein context surrounding each missense mutation from a canonical set of human transcripts in (Hg GRCh37.74). All 9 and 10-mers overlapping the missense mutations were extracted and NetMHCPan(66) was used to predict their affinity to alleles of MHC-I.

### Statistical methods

#### Hypothesis tests

Two-sided Mann-Whitney and Fisher’s exact tests were performed with the 

~~~
R
~~~

 functions 

~~~
wilcox.test
~~~

 and 

~~~
fisher.test
~~~

 respectively. These tests are appropriate as they are non-parametric (distribution-free). One-way ANOVA tests were performed with the 

~~~
R
~~~

 function 

~~~
aov
~~~

 for purity, stromal infiltration, and immune infiltration scores. This test is appropriate as the variance of the scores is similar between the immune infiltration clusters, and ssGSEA scores from 

~~~
gsva
~~~

(56) are approximately normal. P-values were adjusted for multiple hypothesis testing using the 

~~~
R
~~~

 function 

~~~
p.adjust
~~~

 with the “fdr” option.

#### Unsupervised clustering

The unsupervised clustering for tumor samples, immune cell types, genes, and proteins was performed with hierarchical clustering, Ward linkage and Euclidean distance.

#### Random forest prediction of immune infiltration class for SATO patients

A random forest classifier was trained on the TCGA cohort of 415 patients with 10000 trees and otherwise default values in the 

~~~
R
~~~

 package 

~~~
randomForest(67)
~~~

. Training error on the TCGA cohort was 0 percent. This classifier was applied to the ssGSEA scores of the SATO and GERLINGER cohorts to obtain class predictions. The random forest 

~~~
R
~~~

 object and the code to predict the class of a new sample are available upon request.

**Survival analysis**: P-values in Fig. 6b were obtained from univariate Cox proportional-hazards regression models. Kaplan-Meier survival curves in Fig. 6c were plotted for the above-median and below-median equal-size subsamples of the cohort, and a chi-square test statistic for the difference of the curves was computed using a log-rank test.

#### Ratio of cell counts

ssGSEA-based infiltration scores do not follow a discrete count distribution, but are unimodal and approximately normal(56). Therefore ratios of cell counts cannot be determined by simple division of the ssGSEA scores. However, if *a* and *b* represent two cell counts, the log of the ratio *a*/*b* is equal to log(*a*) – log(*b*). Thus, the difference of two ssGSEA scores represents a ratio of cell counts. The CD8^+^ T/Treg and Th17/Th2 ratios in Fig. 6b and 6c denote the numeric difference between the ssGSEA scores for these cell types.

### Code availability

The 

~~~
R
~~~

 code is available upon request.

## Supplementary Materials

**Fig. S1**. Number of genes shared between signatures

**Fig. S2.** Batch effect correction for the three microarray datasets used to derive immune cell gene signatures

**Fig. S3.** Principal component (PC) separation of T cell subpopulations using signature genes

**Fig. S4**. Single gene validation of cell type scores

**Fig. S5**. Pan-cancer analysis of overall immune infiltration score (IIS)

**Fig. S6**. Infiltration scores for selected T cell subpopulations

**Fig. S7**. The signatures where ccRCC is among the highest or lowest across 19 cancer types

**Fig. S8**. Validation of ccRCC immune infiltration classes with the SATO dataset Fig. S9. The correlation between the macrophage and ESTIMATE stromal scores in ccRCC

**Fig. S10**. Grade– and stage–specific APM expression

**Fig. S11**. Prediction of immune infiltration class for Gerlinger *et al.* multiregion tumor samples

**Fig. S12**. Association of immune cell types with clinicopathologic variables Fig. S13. Association of ccRCC immune infiltration classes with genomic alterations

**Table S1**. Gene Signature List

**Table S2.** Single Gene Validation

**Table S3**. Infiltration Score Summary (TCGA ccRCC)

**Table S4**. Significantly Overexpressed Genes in T Cell Enriched Cluster (TCGA ccRCC)

**Table S5**. Significantly Overexpressed Genes in Non-Infiltrated Cluster (TCGA ccRCC)

**Table S6**. Significantly Overexpressed Genes in Heterogeneously Infiltrated Cluster (TCGA ccRCC)

**Table S7a**. Overexpressed gene sets in the T cell enriched cluster (SATO ccRCC)

**Table S7b**. Overexpressed gene sets in the non-infiltrated cluster (SATO ccRCC)

**Table S7c.** Overexpressed gene sets in the heterogeneous cluster (SATO ccRCC)

**Table S8a**. Significantly Overexpressed Genes in TCa Subcluster (TCGA ccRCC)

**Table S8b**. Significantly Overexpressed Genes in TCb Subcluster (TCGA ccRCC)

**Table S9a**. ccRCC APM Mutations (TCGA ccRCC)

**Table S9b.** APM Mutation Statistics (TCGA ccRCC)

**Table S9c**. Expanded List of APM Genes

## Funding

This investigation was supported by: AAH was supported by the MSKCC Department of Surgery Faculty Research Award. AGW was supported by the Stephen P. Hanson Family Fund Fellowship in Kidney Cancer. AAH, AGW, SDK, PR and JAC were supported by the Sidney Kimmel Center for Prostate and Urologic Cancers. RSG was supported by a Medical Scientist Training Program grant from the National Institute of General Medical Sciences of the National Institutes of Health under award number: T32GM007739 to the Weill Cornell/Rockefeller/Sloan-Kettering Tri-Institutional MD-PhD Program. IO was supported in part by the Core Grant (P30 CA008748. YBC was supported by Cycle for Survival of MSKCC. DAS was supported by RO1CA55349 and P01CA23766. SDK was supported by T32CA082088-15. MOL was supported by MSK Translational Kidney Cancer Research Program, and Geoffrey Beene Cancer Research Center. CS and YŞ were supported by NRNB P41GM103504 and GDAC U24CA143840.

## Author contributions

**YS**- concept generation, biostatistical analysis, manuscript writing, manuscript review

**AGW**- concept generation, biostatistical analysis, manuscript writing, manuscript review

**RSG**- concept generation, biostatistical analysis, manuscript writing, manuscript review

**ML**- Multiplex immunofluorescent staining and infiltration calculation

**AL**- biostatistical analysis

**IO-** biostatistical analysis, manuscript review

**NW**- concept generation, biostatistical analysis, manuscript review

**WL**- concept generation, biostatistical analysis, manuscript review

**SDK-** biostatistical analysis, manuscript review

**YBC**- pathologic slide evaluation, manuscript review

**MHV**- concept generation, manuscript review

**JAC**- contributed surgical samples, concept generation, manuscript review

**PR**- contributed surgical samples, concept generation, manuscript review

**VER**- pathologic slide evaluation, manuscript review

**TAC**- concept generation, manuscript review

**EHC**- concept generation, manuscript review

**DAS**- concept generation, manuscript review

**MOL**- Multiplex immunofluorescent staining, concept generation, manuscript review

**JJH**- concept generation, manuscript review, senior advisor

**CS**- concept generation, statistical analysis, manuscript review, senior advisor

**AAH**- concept generation, manuscript review, senior advisor

## Competing interests

None

## Supplementary Figure Legends

**Fig. S1.**
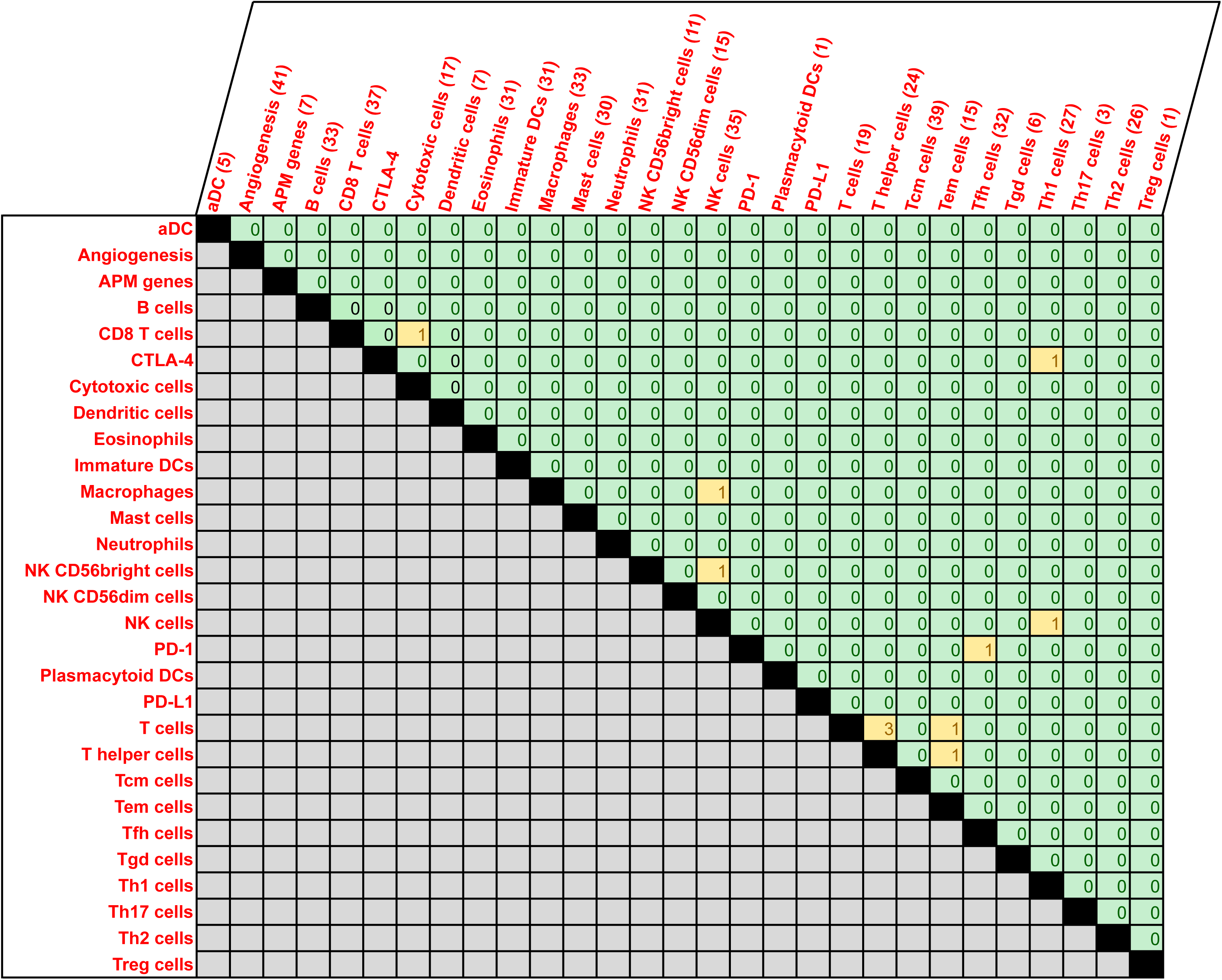
Number of genes shared between signatures. The number in each box denotes the number of genes in common between the corresponding signatures. 98.4% (501) of these genes were used uniquely in only one signature

**Fig. S2.**
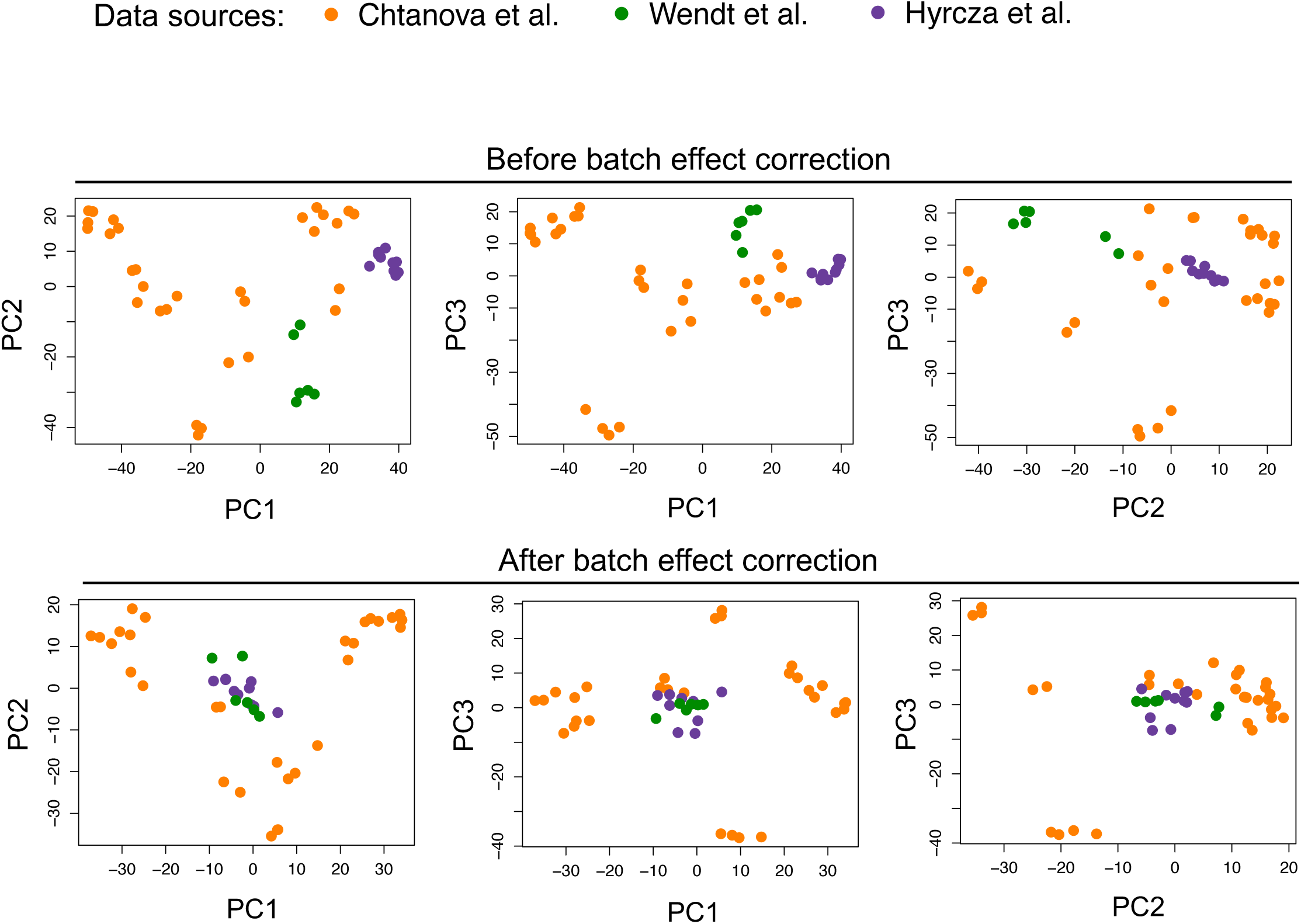
Batch effect correction for the three microarray datasets used to derive immune cell gene signatures. PC analysis on the GCRMA-normalized microarray expression data using 501 signature genes revealed batch effects from the three data sources (top panel). Batch effects were corrected using the nonparametric option in ComBat (bottom panel). After batch-effect correction, cell types of similar lineages but from different data sources clustered together. Cell type labels are given in the batch-effect-corrected PC plot in Fig. 2a.

**Fig. S3.**
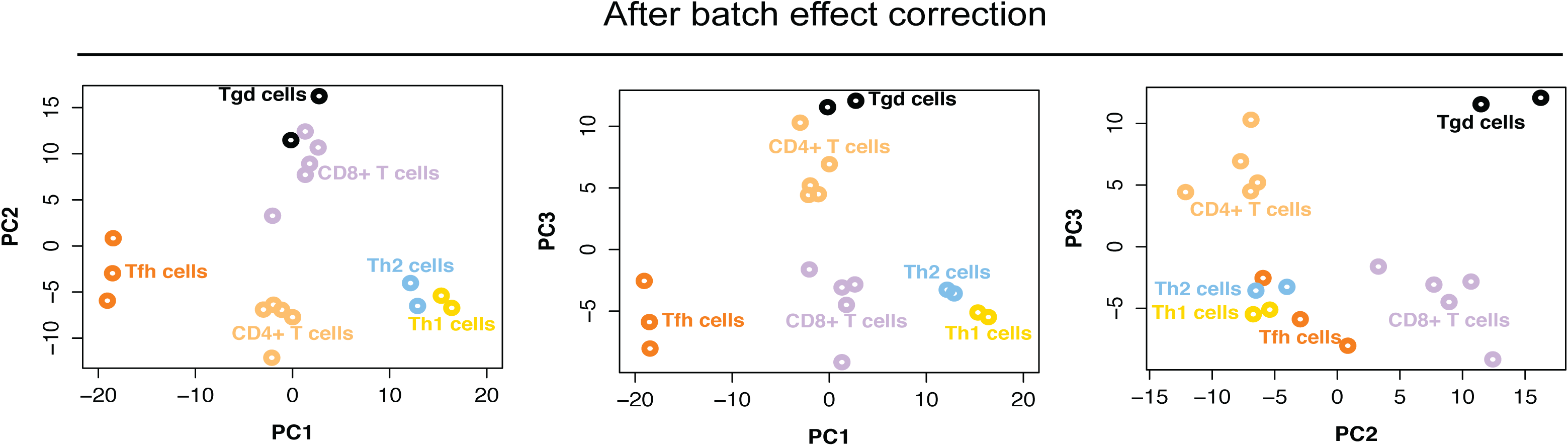
Principal component (PC) separation of T cell subpopulations using signature genes. Microarray gene expression data generated from sorted immune cell types were normalized with GCRMA and filtered to keep only T cell subpopulation samples and signature genes for these subpopulations. Batch effect correction was then performed with ComBat before running PC analysis on the samples. Signature genes achieve a robust separation of T cell subpopulations in this analysis.

**Fig. S4.**
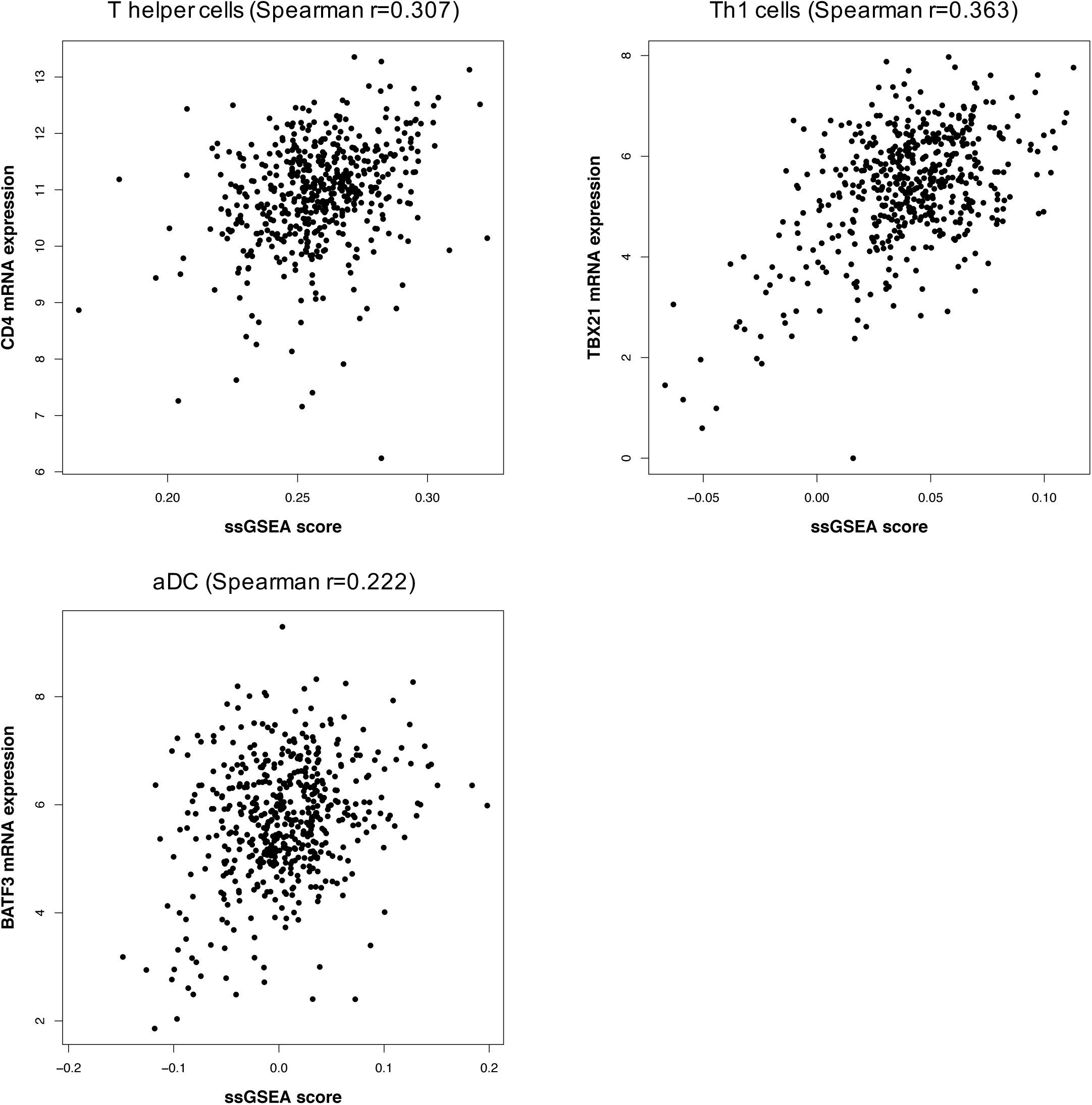
Single gene validation of cell type scores. Spearman correlation between ssGSEA-based immune infiltration scores with the gene expression levels of key immune cell markers for particular cell types. T helper and Th1 cells show good correlations with CD4 and TBX21, respectively. Activated dendritic cells are correlated with BATF3. Gene expression levels are obtained from log2-transformed RNA-seq data.

**Fig. S5.**
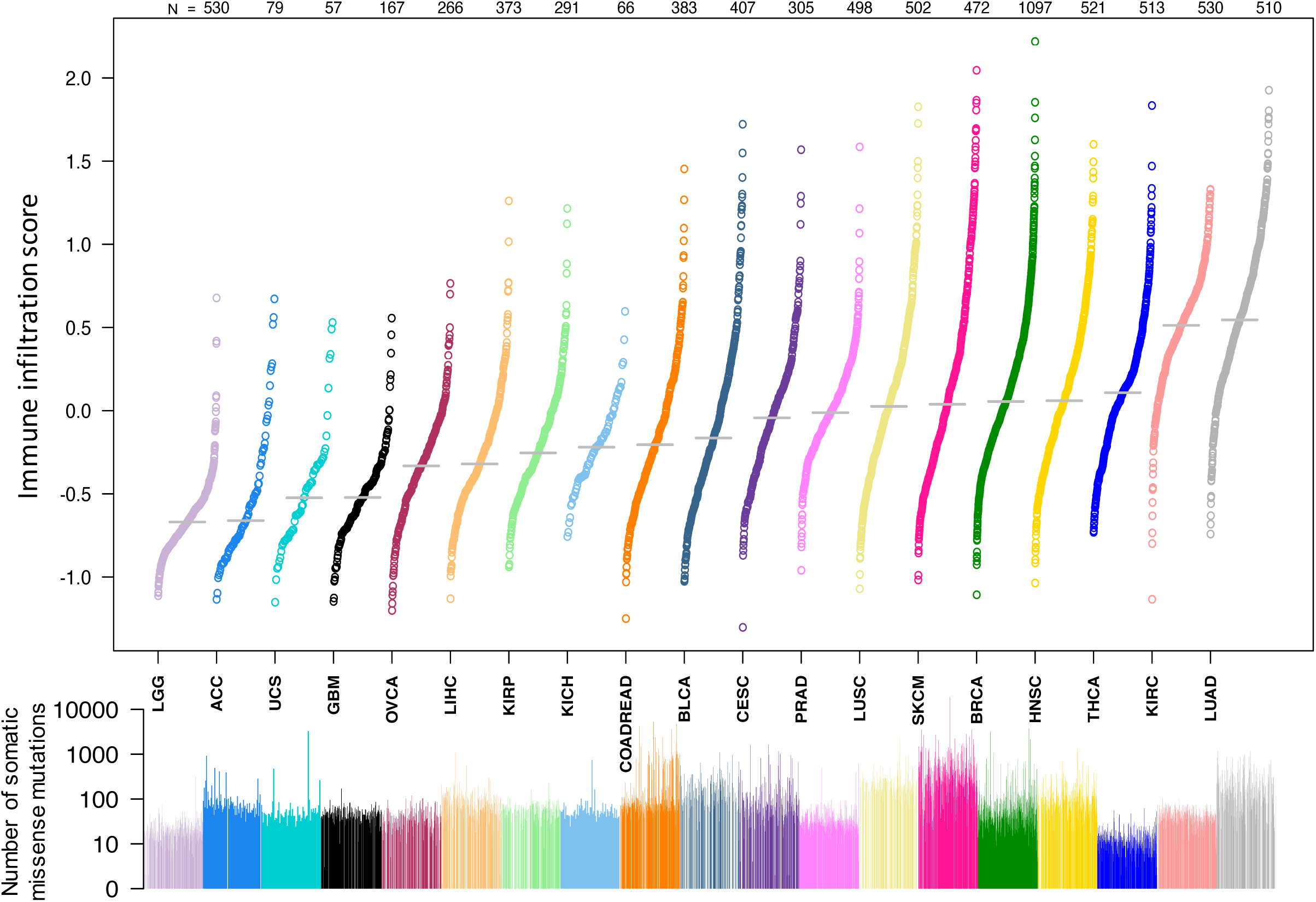
Pan-cancer analysis of overall immune infiltration score (IIS) Overall immune infiltration score (IIS) (top panel) and total number of somatic missense mutations (bottom panel) for 19 tumor types profiled in the TCGA. Each dot represents an individual tumor sample. The order of tumors is largely the same as in Fig. 3a. There is little relationship between immune infiltration and quantity of somatic missense mutations.

**Fig. S6.**
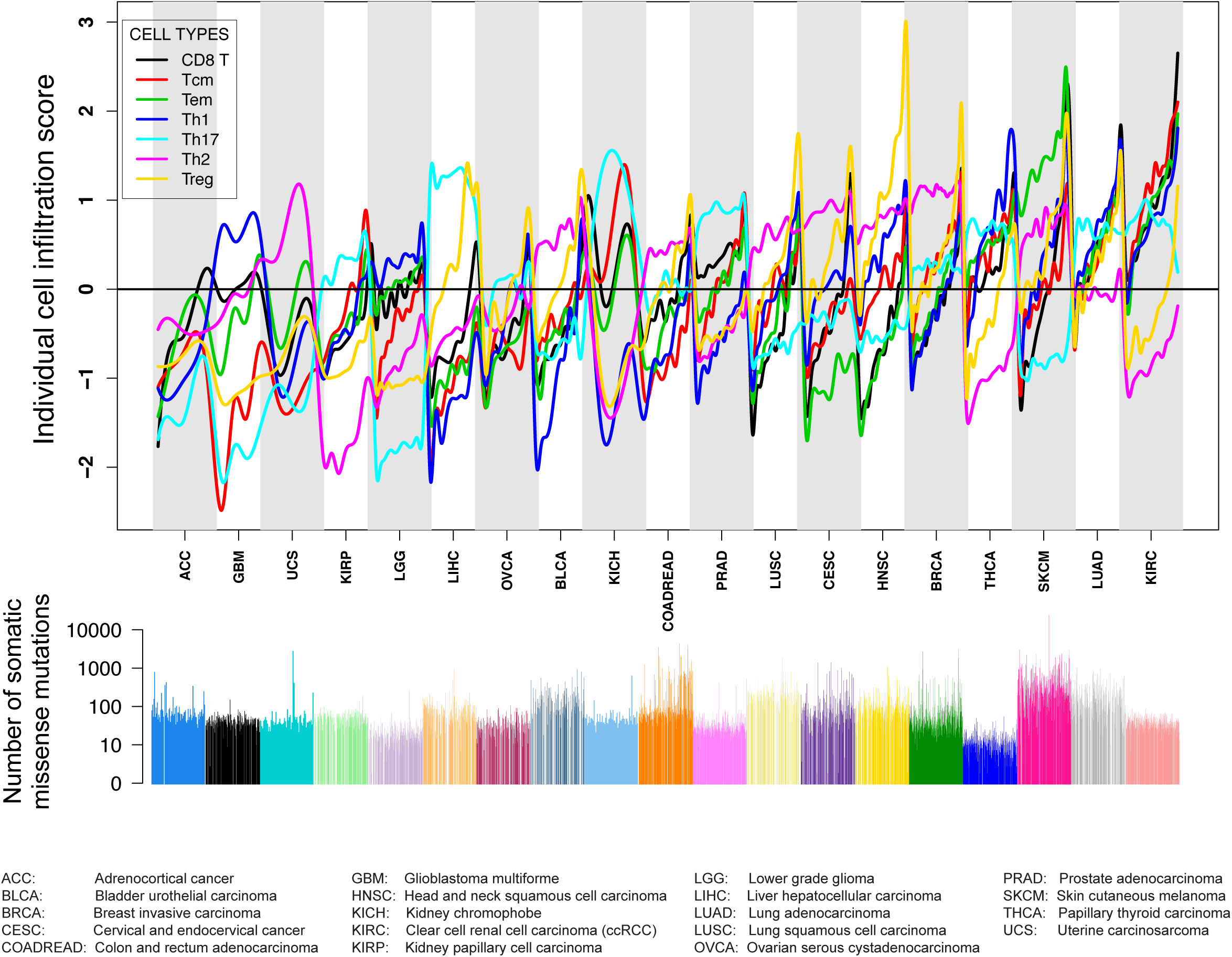
Infiltration scores for selected T cell subpopulations. (Top) Infiltration scores for selected T cell subpopulations (CD8^+^ T, Th1, Th2, Th17, Treg, T effector memory, T central memory cells) that are part of the aggregate TIS and (Bottom) the corresponding mutation load in 19 tumor types in log10 scale. The order of the samples is adapted from the TIS order in **Fig. 3.** A cubic smoothing spline is fit to the data values to generate each curve.

**Fig. S7.**
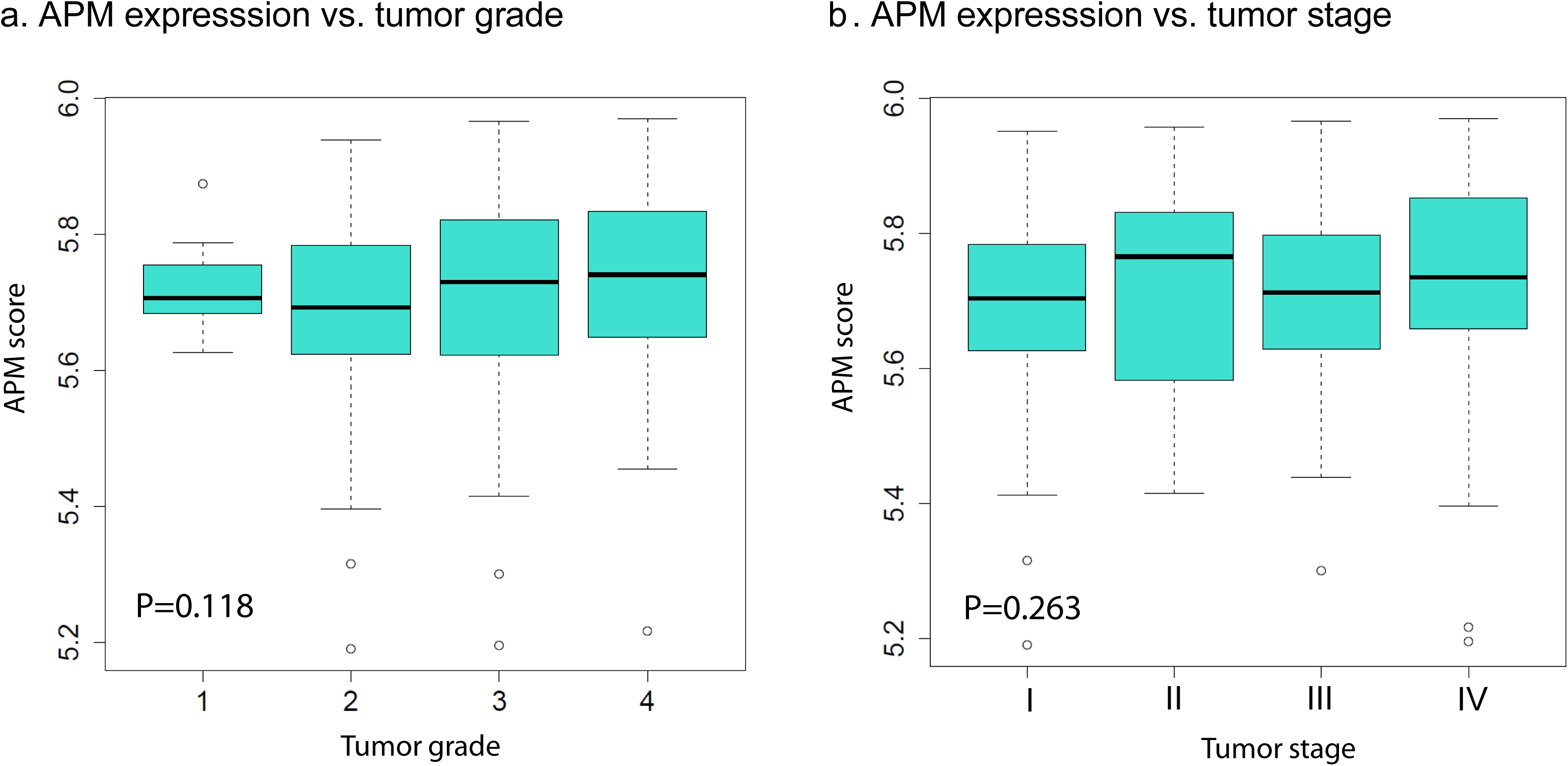
The signatures where ccRCC is among the highest or lowest across 19 cancer types. Analysis of immune cell and angiogenesis levels across 19 human cancers. ccRCC tumors stand out from others by having elevated levels of angiogenesis, several T cell signatures (T cells, CD8+ T cells) along with pDCs, cytotoxic cells and neutrophils. ccRCC tumors are relatively poorly infiltrated by Tregs and Th2 cells.

**Fig. S8.**
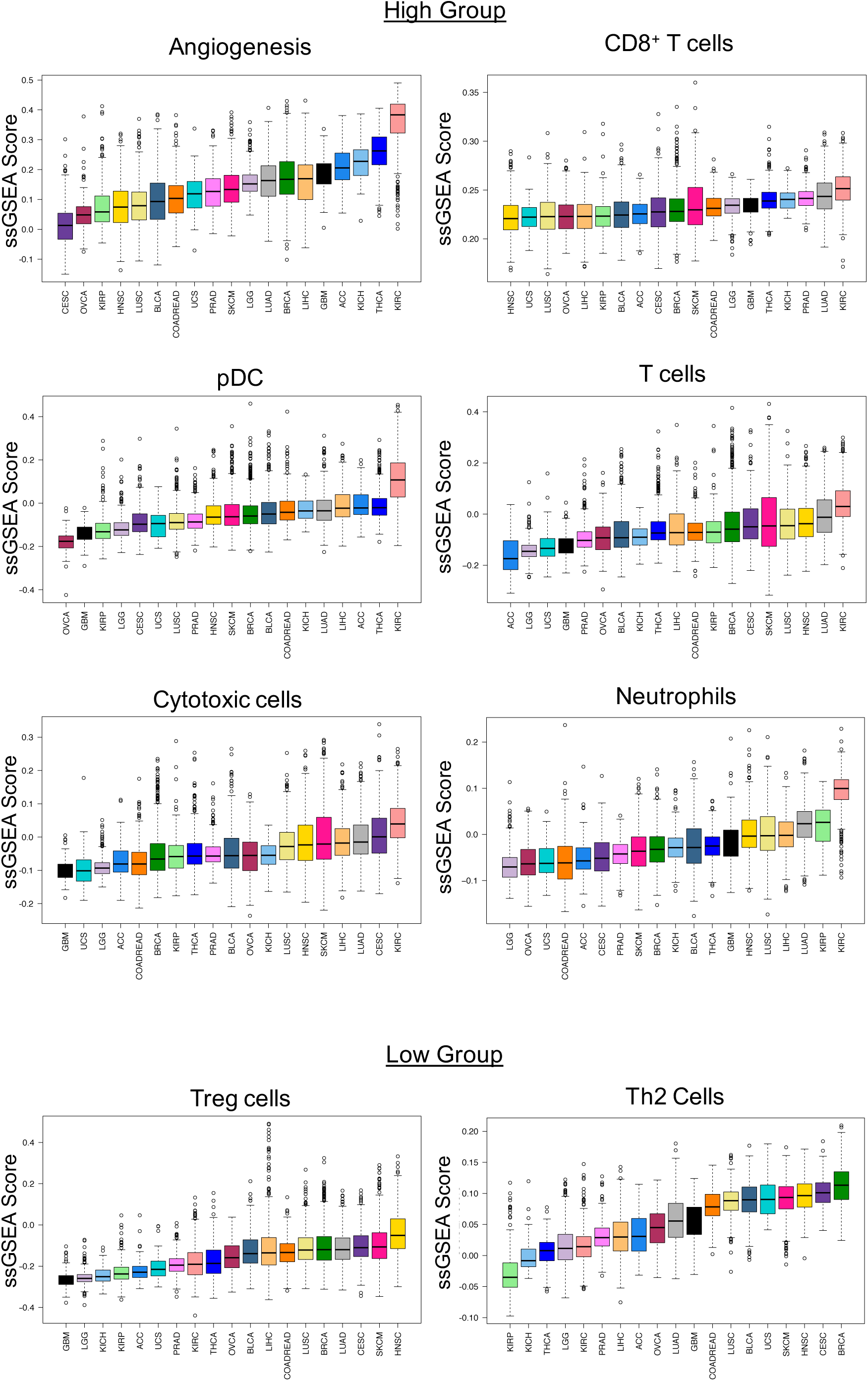
Validation of ccRCC immune infiltration classes with the SATO dataset. (**a**) A random forest classifier trained on the TCGA ccRCC cohort was used to predict the immune infiltration class for 101 patients in the SATO cohort. As was observed in TCGA ccRCC tumors (Fig. 4a), T cell enriched tumors show higher expression of antigen presentation machinery genes, granyzme B and interferon gamma. The order of samples in each class from left to right is by increasing immune infiltration score (IIS). The order along the y-axis is adopted from the TCGA ccRCC heatmap in Fig. 4a. **(b)** Heatmap of genes overexpressed in each immune infiltration class (p-value threshold 0.01). The order along the y-axis is obtained by hierarchical clustering with Euclidean distance and Ward linkage. DAVID gene set enrichment analysis reveals that T cell enriched tumors have overexpression of immune response genes while non-infiltrated tumors have overexpression of mitochondrial genes. These results validate the findings in the TCGA ccRCC cohort.

**Fig. S9.**
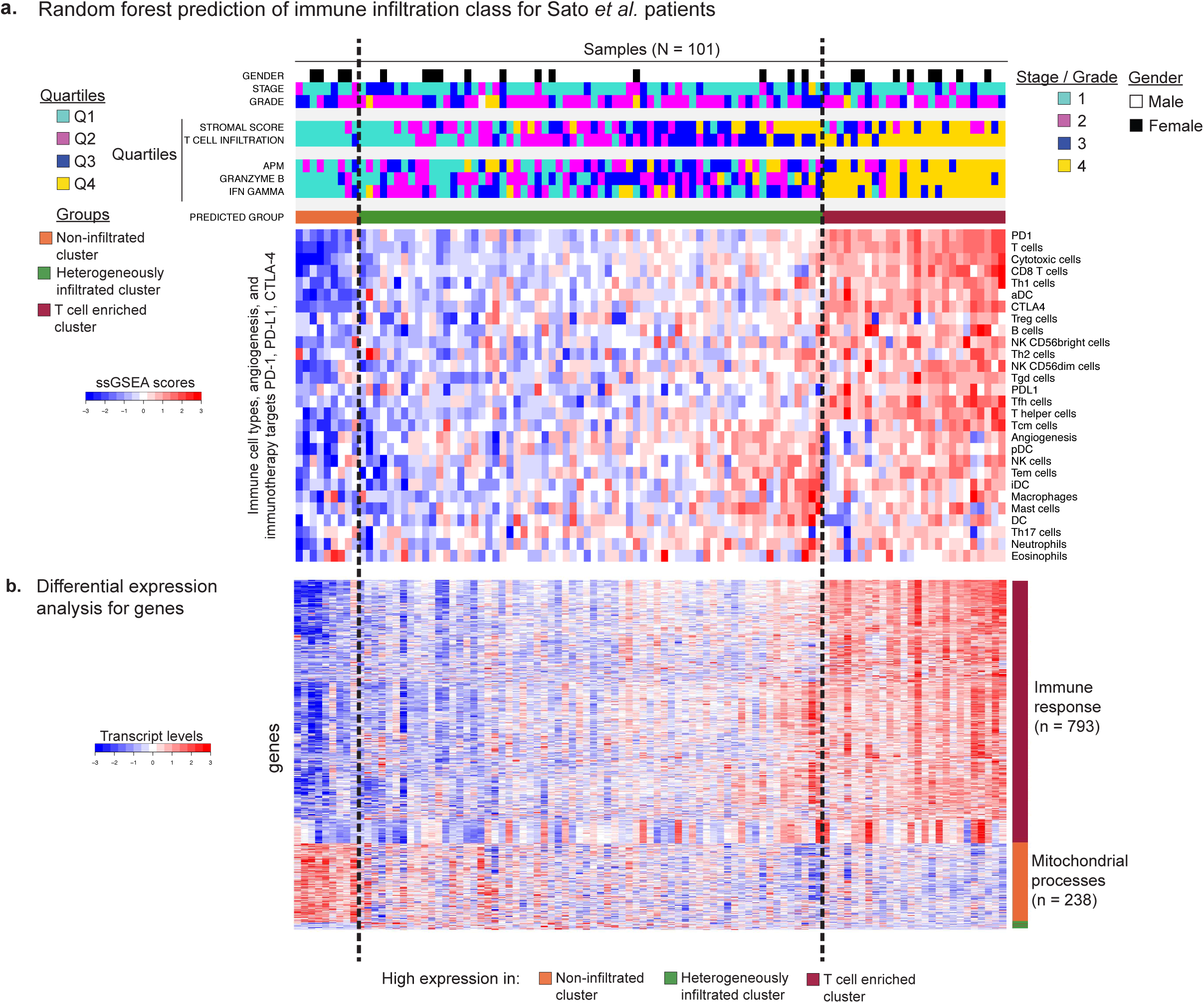
The correlation between the macrophage and ESTIMATE stromal scores in ccRCC. We investigated the association between the macrophage scores in ccRCC and the stromal scores calculated with the gene signature in ESTIMATE. These scores were positively correlated across the entire TCGA ccRCC cohort (Spearman r = 0.561, p < 2×10^−16^).

**Fig. S10.**
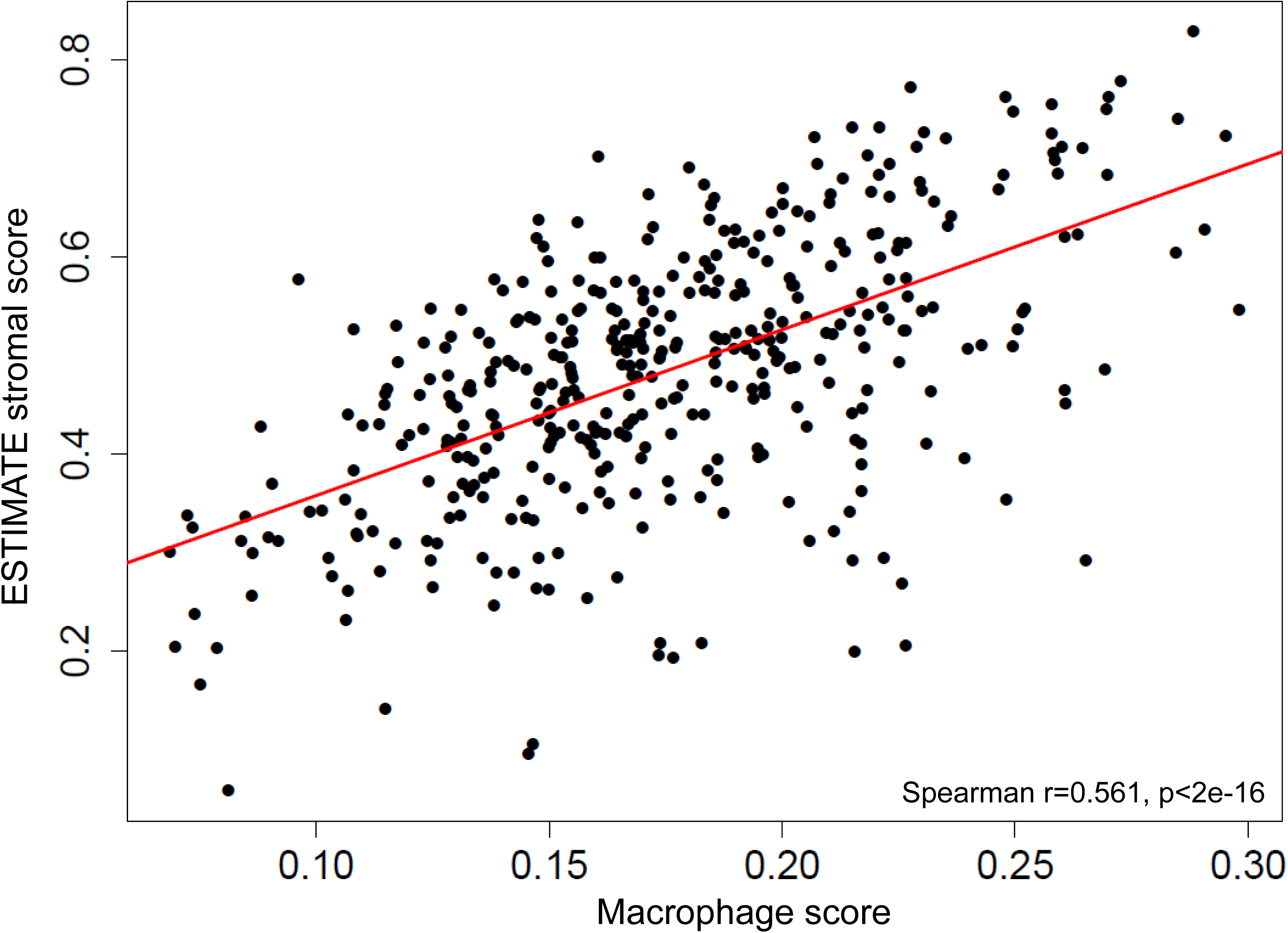
Grade– and stage–specific APM expression. We investigated the association of antigen presentation machinery gene expression with **(a)** tumor grade and **(b)** tumor stage. No significant associations were observed (p = 0.118 and 0.263 respectively, Fisher’s exact test).

**Fig. S11.**
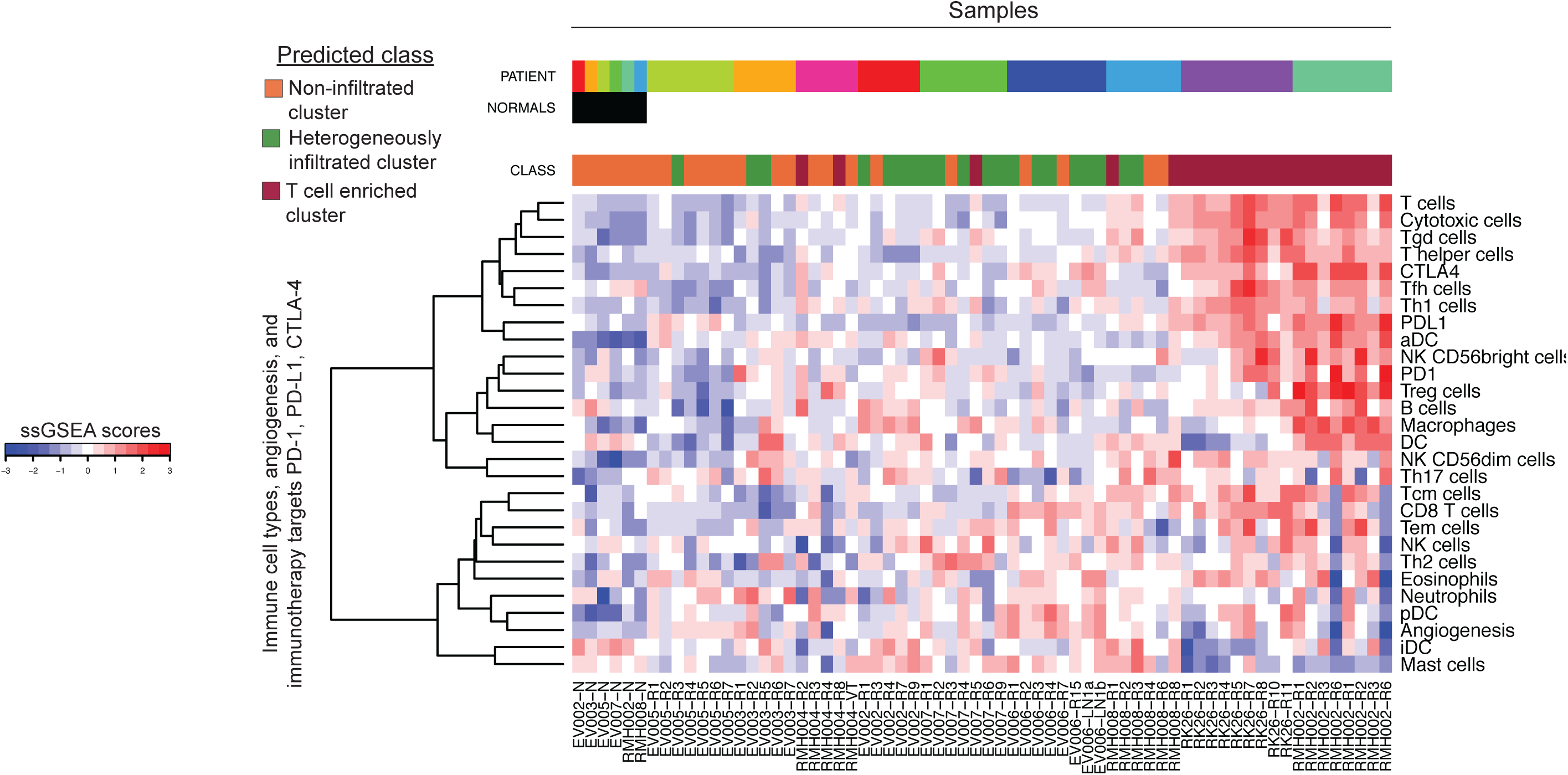
Prediction of immune infiltration class for Gerlinger *et al.* multiregion tumor samples. The immune infiltration class for each sample was predicted with a random forest classifier trained on the TCGA ccRCC cohort. The y axis shows immune cell types and immunotherapy targets ordered according to Ward linkage in hierarchical clustering. The x axis shows the normal samples as a separate group on the left, and the tumor samples from 9 patients. Patients are ordered according to increasing average infiltration level from left to right. Tumor samples within each patient are ordered according to alphabetical order. As can be observed in the results for patients RMH002 and RK26, tumors that exhibit a strong T cell enriched phenotype do not show intratumoral differences and have this phenotype in all sampled regions of the tumor.

**Fig. S12.**
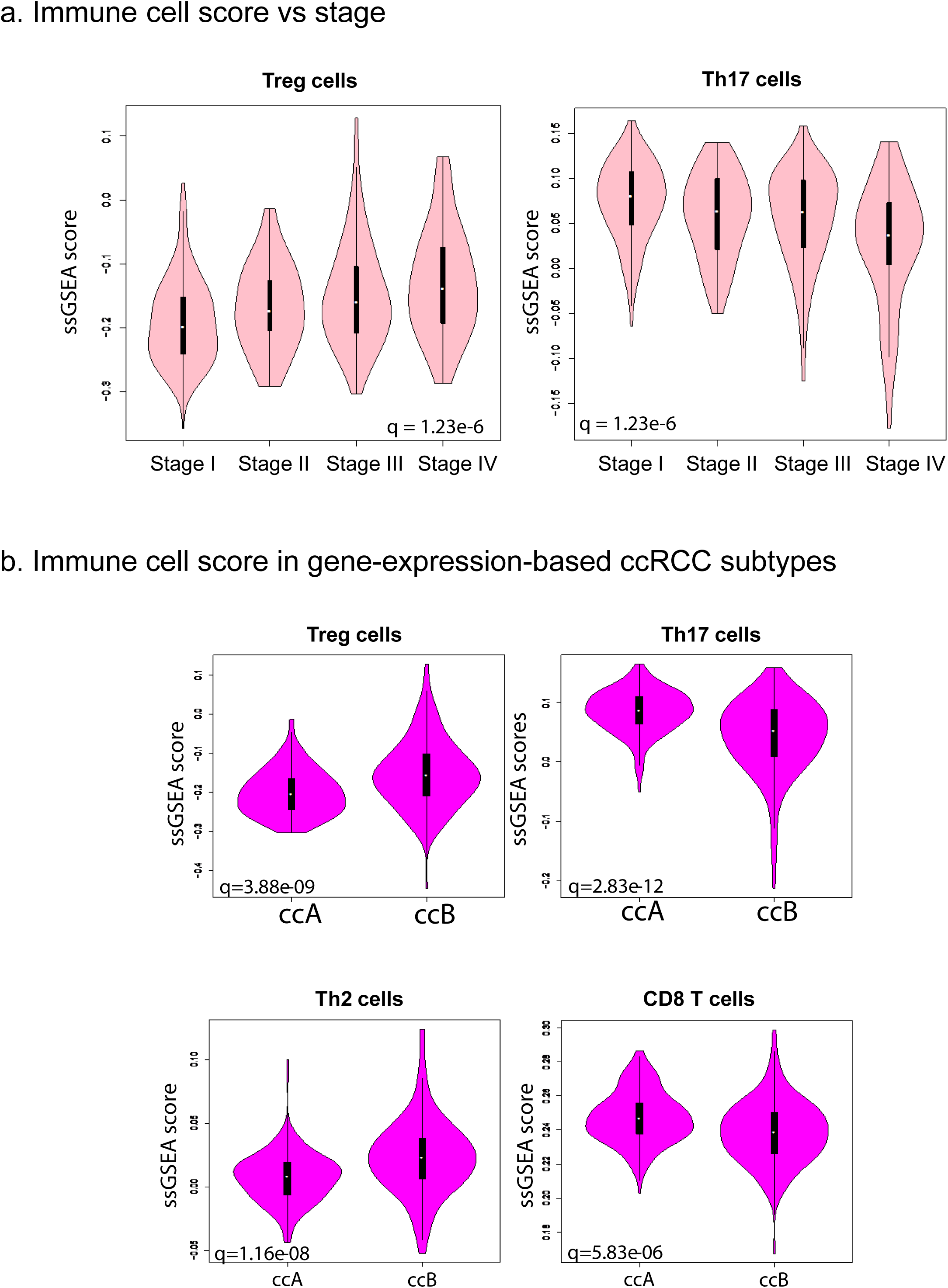
Association of immune cell types with clinicopathologic variables. **(a)** Tumor stage is positively associated with Treg cell infiltration (left panel) and negatively associated with Th17 cells (right panel) in the TCGA ccRCC cohorot. **(b)** Association of immune cell scores with previously defined molecular ccRCC subtypes, ccA and ccB. ccA exhibits significantly higher Th17 and CD8^+^ T cell infiltration levels, but lower scores for Treg and Th2 cells. The former two cell types are associated with improved survival, and the latter two with poor survival (**Fig. 6B**). These findings are consistent with reports that ccA has better prognosis compared with ccB(51).

**Fig. S13.**
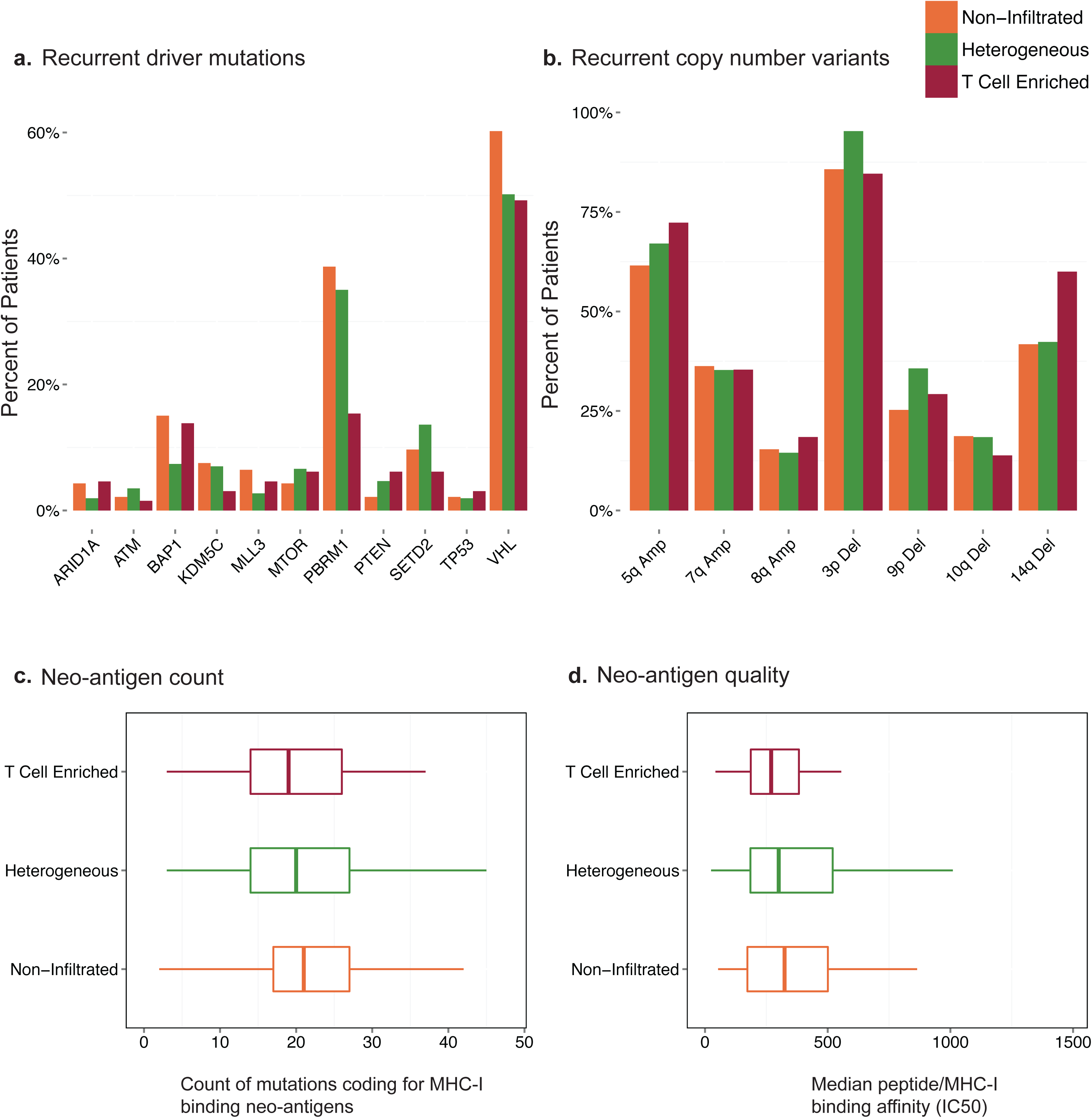
Association of ccRCC immune infiltration classes with genomic alterations. Between the three groups, we found no differences in **(a)** the frequency of recurrent ccRCC driver mutations, **(b)** copy number variants or **(c)** count of mutations that code for at least one neo-antigen predicted to bind to MHC-I (IC50 < 500nM). **(d)** We assessed the overall quality of the neo-antigens found in each cluster by selecting the highest affinity pMHC for each mutation and taking the median of these IC50s (IC50 is inversely related to binding affinity). There was no significant difference in the mutation quality across the three groups of tumors.

